# Acute stress reduces risk-aversion by changing magnitude perception

**DOI:** 10.1101/2025.11.03.685504

**Authors:** Maike F. Renkert, Gilles de Hollander, Gökhan Aydogan, Saurabh Bedi, Paul Forbes, Claus Lamm, Christian C. Ruff

## Abstract

Stress is thought to impair financial decision-making by influencing the willingness to take risks. This effect is commonly attributed to stress-related changes in affective evaluation of rewards and their associated uncertainty. However, existing research has yielded inconsistent findings, and the psychological and neural mechanisms at play remain poorly understood. Here we show that acute psychosocial stress can affect financial risk-taking by altering basic perception of payoff magnitudes, independent of subjective valuation processes. Psychophysical and fMRI modelling reveals that stress reduces risk-aversion without changing choice consistency. A Bayesian perceptual model attributes this effect to an upward shift in prior beliefs about payoff magnitude: Under stress, participants’ choices are guided by relatively more optimistic beliefs about the risky options. A convergent pattern emerges in parietal neural representations of the payoff magnitudes, which show a systematic upward shift in neural coding as a consequence of the stress manipulation. These findings provide converging neural and behavioral evidence that stress influences financial risk-taking by changing neurocomputational processes related to magnitude perception. This has significant implications for psychological and neuroscience theories of stress and risky behavior, with potential applications to policy and clinical interventions.

## Introduction

In modern societies, stress is on the rise, with factors such as economic instability^1^, poor work-life balance^2^, and political or social conflict^3^ exacerbating stress levels. Prolonged exposure to stress has been linked to negative physiological^4^ and psychological^5^ outcomes, such as cardiovascular diseases^6^, anxiety, and depression^7^. Furthermore, stress has also been shown to trigger or exacerbate maladaptive behaviors such as substance abuse^8^ and problem gambling^9^. Outside of the pathological realm, some evidence suggests that stress may negatively affect financial decision-making and could thereby lead to more poverty^10^ and endanger the stability of financial markets^11^. These negative effects of stress might be amplified by the fact that many crucial financial decisions need to be taken during times of substantial change, such as during divorce or unexpected loss of close relatives^12^. Hence, understanding how acute stress changes risky decision-making is important to inform strategies and interventions to mitigate these negative impacts on individuals and society.

Unfortunately, controlled laboratory experiments that have investigated the relationship between stress and financial decision-making have yielded inconclusive results. Some studies report that stress increases risk-seeking^13,14^, others find increased risk-aversion^15,16^, and yet others find no effect at all^17,18^. This apparent inconsistency in results might reflect methodological differences between the studies, relating to how risk attitudes are quantified and how stress is induced experimentally. For example, most of the existing studies have measured risk-taking based on the proportions of risky choices, without considering that this measure reflects changes not only in risk preference but also in choice consistency (i.e., how noisy and, therefore, inconsistent behavior is relative to the expected value difference). Thus, stress-induced changes in risk preferences may have been confounded by, or confused with, the increased noisiness of behavior, as cautioned by methodological studies^19,20^. Additionally, stress is a muti-faceted phenomenon that differs across different elicitation methods, experimental timing schedules, and people. This needs to be taken into account when interpreting previous studies, since these have focused on either acute or more chronic stress^21^, have used different stress elicitation methods that are more physical (e.g., cold-pressor task^22^) or psychosocial (e.g., Trier Social Stress Test^23^) in nature, and have employed different timings of the stressor with respect to the risk elicitation paradigm^24^. Moreover, some studies report that variables like affective valence^25^, arousal^26^ and gender^27,28^ can moderate or diversify the effect of stress on risk-taking, highlighting that these factors need to be accounted for.

Adding to these methodological considerations, the exact neuro-cognitive mechanisms by which stress may influence risk-related behavior remain unclear. For example, one study argued that stress might change risk preferences by promoting habitual/automatic behavior (in light of dual-process theory)^22^, whereas another study suggested that it may do this by impairing cognitive control^29^. Despite this shared focus on disruption of deliberative reasoning processes, these two studies reported opposing behavioral outcomes - increased risk-seeking in the former and increased risk-aversion in the latter. Moreover, while the mechanisms proposed to be changed by stress are interesting and theoretically plausible, they remain interpretations and have not been directly tested with a cognitive model. To our knowledge, no studies to date have directly investigated how stress alters more precisely defined neurocognitive mechanisms (i.e., in terms of a computational cognitive model that can be fitted to data); in particular, no study has so far investigated the impact of stress within a framework that can jointly explain changes in behavior, cognitive processing, and associated neural processes.

Here, we provide such an account of the neurocomputational processes by which stress can change risk-taking in financial behavior – namely, via its effects on the *perception* of the choice problem. Our approach is rooted in recent work specifying how economic decisions can be profoundly influenced by the way we perceive choice-relevant variables^30–33^, and by other lines of work showing that perception can be systematically influenced by bodily and brain states^34–36^. Building on these insights, we investigated how acute stress can, via its effects on perception, alter risk preferences in financial choices. This thorough characterization of the impact of stress on perception and associated economic choices might give us a handle on improving decision-making under stress and could, ultimately, lead to more effective strategies for managing stress in high-stakes environments; for example, by improving the nature and quality of information presentation.

To systematically study the neurocognitive perceptual mechanisms by which stress might alter risky decisions, we employed a novel Bayesian *perceptual psychophysical model*^*37,38*^. This model differs fundamentally from classical economic and finance models. It does not include subjective utility functions—whose curvature typically explains risk aversion—but instead assumes that decision-makers aim to choose the option with the highest objective expected value, without any inherent motivational tendency to avoid or prefer risk. However, critically, decision makers are biased in their perception of the magnitudes at stake due to their noisy perception of the choice variables. Rather than assuming that risk attitudes reflect motivational biases for or against uncertainty^39^, or properties of the subjective valuation of the objective payoff magnitudes^40^, this model mechanistically explains risk attitudes as a natural consequence of the *noisy* and *biased* initial perception of how much money is at stake^30,36^. Note that this perspective differs fundamentally from utility-based theories that simply add random noise to the final choice process^41^, since the latter only model the stochasticity of choices and do not account for any bias resulting from this noise.

Our perceptual model of decision-making under uncertainty is fully consistent with numerous psychophysical and neuroscience studies showing that the brain’s perception of numbers –independent of any subjective valuation – is invariably noisy and thereby fundamentally biased^42,43^. This holds even in the absence of stronger noise (due to the need to infer numerosities from non-symbolic presentations), as is arguably the case with payoffs that are presented as Arab numerals^44,45^. Incorporating these insights from perception research can make microeconomic models more psychologically plausible^30,31,33^, which is particularly important in the context of a phenomenon with clear psychological and physiological consequences, such as acute stress. Moreover, the parameters of our perceptual model are latent variables that correspond to well-defined characteristics of the neurocognitive processes underlying the observed behavior, so that we can directly link them to neural processes and examine how these change during stress.

To do so, we combined the psychophysical model with advanced neuroimaging modeling techniques that allow us to characterize how numbers are represented in the brain - more specifically, in the so-called *parietal approximate number system (ANS)*^*46*^. Populations of neurons in parietal cortex, particularly around the intraparietal sulcus, are known to be tuned to specific numerosities, with activity profiles that are topographically organized and smoothly decrease for neighboring magnitudes ^45,47,48^. These properties make it possible to map numerical representations in humans using functional MRI together with numerical population receptive field (nPRF) modeling^48,49^, which estimates the preferred numerosity and tuning of cortical responses. Importantly, the fidelity of these representations has been shown to predict individual differences in perceptual acuity and decision-making^33,36,50^. We employed this combined cognitive and nPRF modeling approach to characterize how stress changes decision-making under risk, via its influence on Bayesian inference about the payoffs relevant for choice, as extracted from the noisy parietal population code of magnitudes. Our cognitive modeling framework proposes that risk aversion is a consequence of Bayesian perception, with a central tendency effect^31,36^ that directly relates in its strength to the degree of noise in perception/representation of the payoff (likelihood). Specifically, the posterior estimate of the perceived payoff magnitude (on which the decision/comparison is ultimately based) is pulled towards prior beliefs about which potential payoffs are plausible in the first place. The resulting underestimation will –independent of the exact prior beliefs– be inherently stronger for the larger risky option than for the smaller safe payoffs^30^, making the safe option appear to be relatively more attractive and thus making decision-makers, on average, more risk-averse (illustrated in Figure 3A). Using this framework, our previous studies have already established that this perceptual bias is linked to noise in parietal magnitude representations, which can comprehensively account for interindividual and situational differences in risk aversion^33,36^. Here, this modeling framework enables us to disentangle the different ways by which stress might interfere with the perceptual process that results in overt risk preferences.

More specifically, stress may affect neurocognitive processing in two ways that we can differentiate using our model: Decision-makers could either become more or less noisy in the perception and processing of the relevant payoff magnitudes, or they could change their expectations and thus prior beliefs about potential payoffs. The former, a change in noisiness, would lead to decreased precision with which the decision maker represents the option payoffs. This would decrease choice consistency (the relationship between the expected value of choice options and choice proportions) and lead to larger central tendency effects, expressed as either more pronounced risk seeking or stronger risk aversion, depending on the size of the payout relative to the range of payouts that the decision-maker’s believes are most plausible^36^. The latter, a change in prior beliefs about the plausible range of the offer values, would be reflected in a shift in the indifference point, which uniquely characterizes changes in risk preferences independently of the noise in cognitive representation and response behavior^30,51^.

Importantly, when only looking at choice proportions, these effects cannot be separated from changes in any factors that make choices less consistent (e.g. attentional lapses) with an unchanged risk preference and/or payoff perception. Therefore, it is crucial to model the data in a way that quantifies how cognitive noise and prior expectations (not just choice consistency) changes under stress^52,53^. Both these possible effects appear plausible in light of earlier studies on the influence of stress on decision making in a broader sense. On the one hand, some studies have found stress to induce more noise in decision making processes^54,55^ while others have reported improved attention and thus more consistent behavior upon stress exposure^56^. On the other hand, several studies have also reported both increased^22^ and decreased^57^ behavioral biases under stress. Here, we were able to test how changes in both these mechanisms may account for altered risk-taking under stress, within one single neuro-computational framework: In both our Bayesian cognitive model and the neural nPRF model based on fMRI data, we could measure whether neural magnitude representations become noisier and/or reflect a shift in average reference point as a function of stress. This integrated test of competing hypotheses, in both behavioral and neural data, thus offers more mechanistic insights than modeling approaches that are agnostic to the underlying neuro-cognitive processes^58^.

Using our cognitive and neurocomputational model-based approach, we find that acute stress affects risk preferences without changing choice consistency. Generally, most subjects start from a risk averse baseline, but only for the stressed participants, this preference shifts towards risk neutrality in the second session, as subjects become significantly more risk-seeking. Intriguingly, we show that the neural code for presented payoff magnitudes in parietal cortex shifts in parallel. Thus, the cognitive and neural modeling results align in their mechanistic explanation: Both suggest that prior beliefs about payoff magnitudes become, on average, more optimistic, so that under stress, the risky option is perceived as more attractive and is therefore chosen more frequently.

## Results

### Study design, risk task, and stress manipulation

To study the effects of stress on financial decision-making under uncertainty, we opted for a mixed design with repeated (pre- and post-) measures within each experimental group. This design is particularly powerful for measures that are highly variable but relatively stable across repeated measures within single individuals, such as risk preferences^59^. Subjects came to the lab twice. In the first session, we obtained a baseline measure of risk preference; in the second session, we repeated this measurement during an experimental stress manipulation or a matched no-stress control manipulation (Figure 1A). We could thus model the effects of stress as a session × group interaction while accounting for individual differences through subject-level random effects.

**Figure 1.**
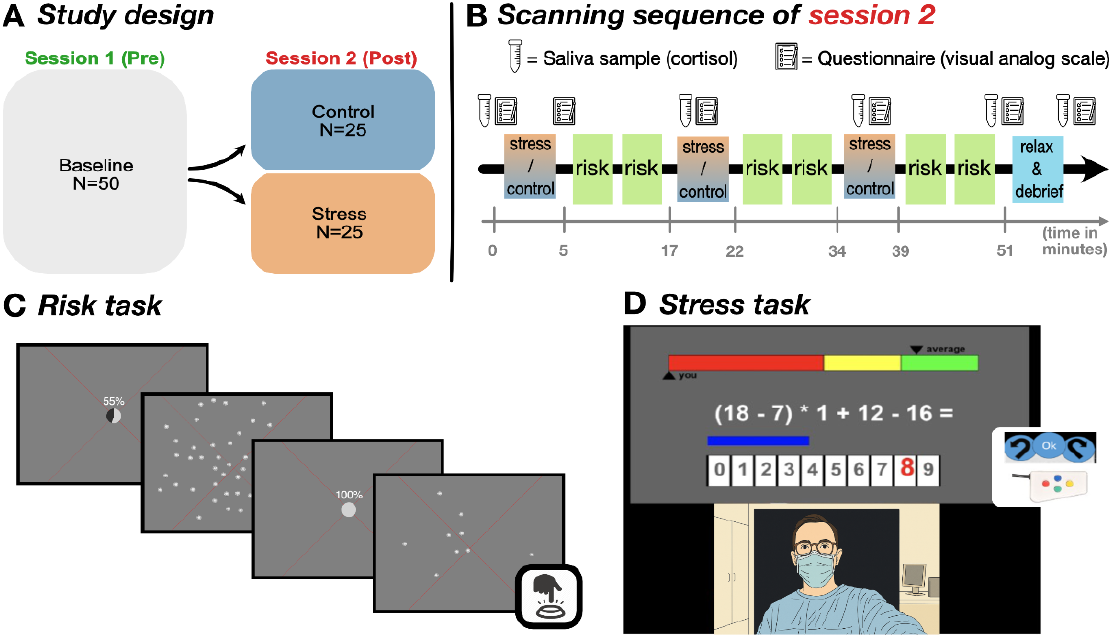
Tasks and study design illustrations. **A. Illustration of repeated-measures design**. In the first (baseline) session, all participants (n=50) performed the risk task. In the second session, the risk task was interleaved with either a control (simple calculations) or a stress (calculations under time pressure plus deceptive negative performance feedback) task. Subjects were equally distributed across those two conditions, controlling for the baseline values of the measures of interest (n=25 each). **B. Sequence of tasks and sample collections in session 2**. At the indicated time points, saliva samples for cortisol measures and emotional state questionnaires were acquired. **C. Illustration of risky choice task**. Two choice options were presented sequentially. Subjects were first shown a pie chart and percentage that indicated if the upcoming option was the risky or safe option. Then, payoff magnitudes were presented as a cloud of coins. As soon as the second payoff was presented, subjects were allowed to indicate their choice using the button box. **D. Illustration of stress manipulation**. Adapted version of the Montreal Imaging Stress Test (MIST)^64^. Participants were presented with complicated arithmetic problems they had to solve under time pressure. The difficulty of the presented problems was adapted dynamically so as to be slightly too hard for the participant to solve in time. A blue bar presented below the problem indicated the time remaining, and the bar on top represented the adaptive current performance level. The bottom part of the screen displayed a live video feed of the experimenters who were critically observing the participant and who asked the participant to make more of an effort as the data would not be usable otherwise. Control participants did a similar task but had no time restriction, no performance feedback, and no negative social evaluation via video..

For the baseline session, all participants performed a series of incentive-compatible risky choices in which they had to repeatedly choose between a safe (certain payout) and a risky option (with a payout probability of p=0.55) that both varied in payoff magnitudes across trials^30,33^. Following our previous work^36^, the varying payoff magnitudes were presented as stimulus arrays consisting of 1-CHF coins. On every trial, the risky and safe options were presented sequentially, with this order being counterbalanced across trials. Participants indicated their choice upon presentation of the second option, requiring them to retain the first option in working memory. As we have shown in earlier work, this retention introduces noise into the representation of the first option (Figure 1C)^36^. For the second session, the risky choice task runs were interspersed (Figure 1B) with either an adapted version of the well-established Montreal Imaging Stress Induction Task (MIST^60^ - Figure 1D) with additional social evaluative threat or a simple control calculation task without any social evaluative (or other) threat components.

Confirming the effectiveness of our stress manipulation, only the MIST but not the control manipulation induced a strong stress response: At multiple time points following the stress manipulation, we found reliable increases in cortisol levels (comparison of area-under-the-curve (AUC) values between stress and control (t(50)= -5.35, p = 2.55*10^-6^; see methods for details) and self-reported stress levels (within-subject mean stress rating from the questionnaire (t(50)= -4.55, p = 3.78*10^-05^; see Figure S1 for a plot of the distributions of the subjectwise physiological and psychological stress levels). While cortisol is a key marker of stress along the hypothalamic-pituitary-adrenal (HPA) axis, stress also modulates autonomic nervous system (ANS) activity. To assess potential shifts in the balance of sympathetic and parasympathetic activity following the stress manipulation, we also analyzed heart rate variability (HRV), a marker of sympathetic and parasympathetic tone^61,62^. Supporting the success of our stress manipulation, the low-frequency/high-frequency HRV-ratio, a proxy for sympathetic/parasympathetic balance^62,63^, significantly increased more in the stress compared to the control group (U(50)=203, p=0.034; Mann-Whitney test). Moreover, we found numerically stronger decreases in the stress group in the Standard Deviation of Normal-to-Normal intervals (SDNN, t(50)=1.48, p=0.15) and Root Mean Square of Successive Differences (RMSSD, t(50)=1.47, p=0.15); although these effects were not statistically significant, they were in the expected direction for parasympathetic withdrawal under stress, as indicated by a recent meta-analysis on the effect of acute stress on HRV measures^62^ (see Figure S2 for a comprehensive illustration of all heart-rate related measures tested between groups for individual sessions and session differences).

### Acute stress induces more risk-seeking

As an initial test for overall stress effects on risk preferences and the underlying decision-making mechanisms, we fitted an integrated psychophysical probit model to each participant’s behavioral data, with session and group as regressors (Figure 2A). This standard psychophysical model describes how the probability of choosing a risky option increases with the relative attractiveness of the risky option compared to the safe option (i.e., the log-ratio of risky versus safe payoff), quantifying the consistency of participants’ responses. Critically, it also yields an estimate of the point-of-indifference, a measure of risk preference^30^ that captures the ratio at which participants are indifferent between risky and safe options (i.e., choose the risky option 50% of the time). Following earlier studies^30,33,36^, we quantified the point-of-indifference using the risk-neutral probability (RNP), i.e., the hypothetical probability associated with the presented risky payoff that would make a risk-neutral decision-maker show the same point-of-indifference as the participant. Risk seeking (or risk-aversion) is thus reflected in RNPs above (below) 55%, and the degree of risk/seeking or risk aversion is quantified by the absolute distance of the RNP from 55%. Although both groups became slightly more consistent in their choices from session 1 to session 2 (mean slope = 2.035, 95% Credible Interval (CI)=[1.559, 2.531]; average main effect of session on the slope=0.276, 95%CI=[-0.023, 0.59], *p*_*bayesian*_ = 0.037), this effect did not interact with the stress manipulation (mean=-0.05, 95%CI=[-0.504, 0.377], *p*_*bayesian*_ = p=0.418), revealing that the acute stress manipulation had no effect on choice consistency. Conversely, the stress manipulation did lead to increased risk-seeking (i.e., it moved the indifference points from general risk aversion towards more risk-neutral behavior) without changing the consistency of the responses. Specifically, participants became more risk-seeking –and thus closer to risk-neutral– in the experimental group (RNP increased from 47.7%; CI =[38.1, 57.6] to 55.6%, 95%CI =[42.2, 69.3], *p*_*bayesian*_=0.003) but not in the control group (from 49.7%(CI =[40.1, 58.9] to 50.9% (95%CI =[40.2, 62.7]))], *p*_*bayesian*_=0.298). The interaction group x session was also significant (mean=0.07, 95%CI =[-0.01, 0.14], *p* _*bayesian*_ =0.034; Figure 2B).

**Figure 2.**
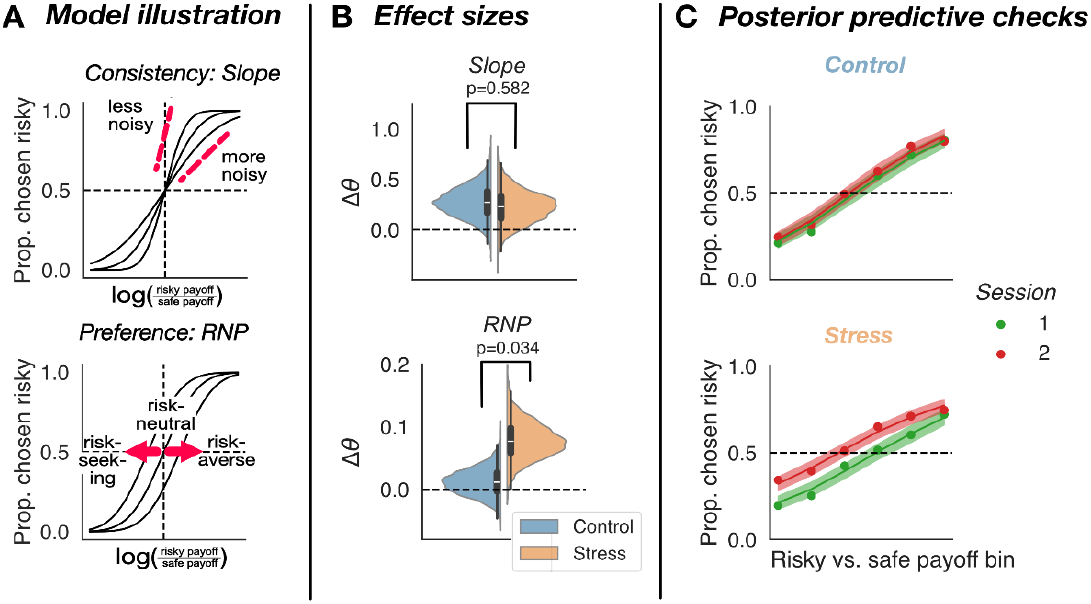
Stress alters observed risk preferences without changing choice consistency. **A. Illustration of possible stress effect on risk-taking with hypothetical psychophysical curves**. Effects on choice consistency (randomness) would be reflected in changes of the slope, whereas effects on preference would be reflected in changes in the point-of-indifference (i.e., the ratio between risky and safe payoffs where the participant chooses both with equal probability). This indifference point can be elegantly quantified by the Risk Neutral Probability (RNP). The more risk-seeking participants are, the higher their risk-neutral probabilities will be. **B. Posterior estimates of effect sizes**. In the stress group, we find evidence for an increase from first to second session (stress effect) on RNP but not slope. No such effect is evident in the control group, and the increase is stronger in the stress than the control group. **C. Posterior predictive checks (PPCs) of the fitted model**. The model captures that only the average choice curve of the stress group, but not the control group, shifts leftwards, implying more risk-seeking behavior. Binned average response proportions always overlapped with the 95% credible interval of the PPCs (shaded area).

### Acute stress leads to more optimistic beliefs about magnitudes

To identify the neurocognitive mechanisms by which stress affects risky choice, we used a computational cognitive model that frames risky choice as a perceptual Bayesian inference problem^36^. In line with the estimated psychophysical curves, the fitted model showed that acute stress did not affect neurocognitive noise. Rather, it modified the prior beliefs of the participants about potential payoff magnitudes. As a result, participants did not become more (in)consistent in their choices but shifted their indifference points to more risk-seeking behavior.

In our perceptual model of risky choice (perceptual memory-based choice model - PMC)^30,36^, the prior beliefs about the plausible range of the payoff magnitudes are combined with the incoming evidence of the payoff at stake, weighted by their respective precision (inverse of the noisiness), to derive and compare the percepts (i.e., the decision maker’s mean posterior estimates) of both payoffs. This model enables us to decompose behavior into multiple latent processes: On the one hand, the precision of the perceptual evidence (separate for the first and secondly presented option, accounting for perceptual noise in both options as well as additional working memory noise for the first); on the other hand, the mean and dispersion of the participant’s prior belief about possible payoff magnitudes (i.e., what payoffs people expect, and how certain they are about these expectations, see Figure 3A and methods for more detail). We allowed session, group, and their interaction to affect both the noisiness of the evidence as well as the mean of the priors, to disentangle potential stress effects on different kinds of cognitive noises (working memory and number sense precision) as well as biases due to expectations (prior beliefs of payoff magnitudes). Note that we have shown in earlier work that all these parameters can reliably be recovered with the number of trials we used here ^65^. Our model-based analysis showed that stress had a specific effect on the means of the priors (interaction between group and session, mean=0.51, 95%CI=[0.178, 0.852], *p*_*bayesian*_ =0.002): Subjects in the stress group had a more optimistic belief about potential payoffs in the second compared to the first session (*p*_*bayesian*_ =0.009, mean=0.388, 95%CI=[0.084, 0.668]), whereas no such effect was evident in the control group (*p*_*bayesian*_ =0.897, mean=-0.120, 95%CI=[-0.302, 0.07]; Figure 3C - see Table S1 for posterior distributions of each parameter for each group at each session). The widths of the priors were not affected by either session or stress, suggesting that these factors did not change the way in which people weighted risky/safe magnitude information differently (statistics in Table S2). Furthermore, stress did not specifically affect the noisiness of the evidence for the two options (first option: *p*_*bayesian*_ =0.259, mean=0.074, 95%CI=[-0.157, 0.299]; second option: *p*_*bayesian*_=0.596, mean=-0.026, 95%CI=[-0.248, 0.197]); however, we found an overall session effect, for both groups, on the noisiness of the representation of the first option (main effects of session, *p*_*bayesian*_ =0.045, mean=-0.128, 95%CI=[-0.276, 0.022]), but not the second option (*p*_*bayesian*_=0.335, mean=-0.030, 95%CI=[-0.174, 0.111]). This is again consistent with the results of the psychophysical model, but shows that the increase in choice consistency (and decrease in noise) over sessions mostly reflects a practice-related increase in working memory precision for the item presented first.

**Figure 3.**
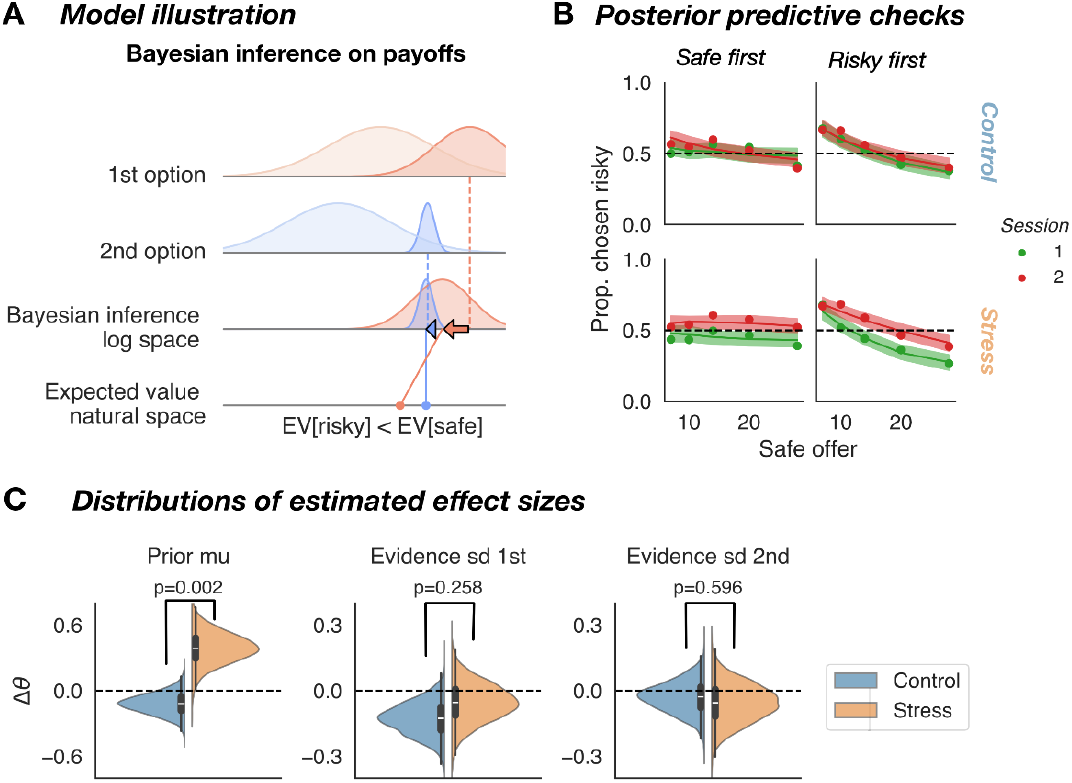
A. The perceptual memory-based choice model (PMC^36^) of risky choices as Bayesian inference process. The model assumes that participants employ Bayesian inference on noisy neurocognitive representations of the objective payoff magnitudes. In this example, the likelihood of the first-presented (risky) option is more dispersed (dark red distribution; top row) than that of the second-presented (safe) option (dark blue distribution; second row), because it needs to be stored in working memory. Both likelihoods are combined with their respective priors (light-colored distributions) to yield the final percept (posterior estimate). For large payoffs (high stakes trial), as in this example, the priors lie to the left of their corresponding likelihoods, resulting in underestimation (the leftwards shift of the posteriors - dark colored distributions in the third row) that is proportional to the noisiness of the evidence. The difference in the degrees of underestimation can make the expected value (fourth row) of the risky option appear to be smaller than that of the safe option. **B. The model can account for order and range effects.** The proportion of risky choices varies systematically from small to high-stakes trials. Critically, whether the proportion of risky choices increases or decreases as a function of stake size depends on which choice option is presented first (i.e., the safe or the risky option). This pattern can be accommodated by the model, as shown by the posterior predictions (lines with shaded areas around them - 95% CI). Only for the stress group, this pattern shifts upwards (more risky choices on average), but it does not change the linear relationship between overall stake size and the proportion of risky choices. **C. Posterior distributions of model parameters**. Only the mean of the prior (Prior mu) shows a shift from first to second session that is significantly larger for the stress than for the control group (see Table S1 for posterior distributions of each parameter for each group at each session). This shows that stress affects magnitude perception mainly by changing payoff expectations.

Crucially, our model was well able to capture the complex pattern of stake and order effects evident in our data (Figure 3B; see also^38^): Subjects were, on average, risk-seeking for small stakes but became risk averse for large stakes. Furthermore, this tendency increased when the risky option came first (and therefore was represented more noisily and thus underestimated more strongly). In line with the general assumptions of our model, both groups showed this pattern consistently across both sessions; however, only for the stress group did the curves shift upwards from the first to the second session, in line with the increased risk-seeking captured by the stress-induced shift in the prior.

We used formal model comparison techniques to quantitatively evaluate our PMC model against an established Expected Utility model^39,40^, as well as to assess whether incorporating an effect of stress improved model fit to the data. Specifically, we used expected Log-Predictive Density (ELPD), a state-of-the-art model comparison metric^66^, to compare four different models (see Figure S3). The PMC models vastly outperformed the EU models and the best-fitting model out-of-four was a PMC model that allowed for an effect of stress (specifically, a group x session interaction). An EU model that allowed for an effect of stress also performed better than a baseline model. The PMC model including a stress effect had the highest ELPD (−6083.57) and a stacking weight of 99.9%. The next-best model was the PMC model without a stress effect (ELPD = −6144.87, dSE = 10.21; stacking weight <0.001%). The third-ranked model was the EU model with a stress effect (ELPD = −6871.06, dSE = 37.83; stacking weight <0.001%), and the worst-performing model was the EU model without a stress effect (ELPD = −6872.95, dSE = 38.19; stacking weight = 0.0018%).

### Stress leads to upward shift in neural magnitude coding in parietal cortex

In addition to the model-based behavioral analyses of the perceptual mechanisms that alter risk preferences under stress, we examined the neural mechanisms underlying altered magnitude perception under stress. The neural evidence aligned well with the results of the cognitive modeling: Acute stress led to an upward bias in neural number processing, without affecting the noisiness of the corresponding representations.

We investigated neural payoff-magnitude processing with an inverted numerical population receptive field (nPRF) model. This encoding/decoding-approach (see also^33,36^) consisted of three steps: First, we fitted numerical receptive field models to each voxel *i*, based on single-trial brain activity patterns *y* _*i*_: *f*_*i*_ (*n*, θ_*i*_), where the parameter vector θ_*i*_ consists of a preferred numerosity, a dispersion parameter, as well as an amplitude and baseline activity parameter. Second, we also fitted a noise model to the most responsive voxels in the parietal cortex, defined as a multivariate t-distribution with the covariance matrix of the residuals of *f*_*i*_ (*n*), Σ. The combined encoding and noise model allowed us to define a likelihood function for bold activity *a*, given a presented numerosity *n, p*(*a*|*n*). In the final test phase, we inverted the encoding model to derive a posterior over numerosities, given the brain activity of a specific trial from an unseen test data set (a fMRI run that was held out of the training set), *p*(*n*|*a*). This inverted model thus allowed us to get a mean posterior *E*[*n*|*a*], which is the ‘neurally decoded’ numerosity for a given trial. Figure 4A provides an example illustration of parietal nPRFs as cortical patches that show tuning to specific numerosity.

**Figure 4.**
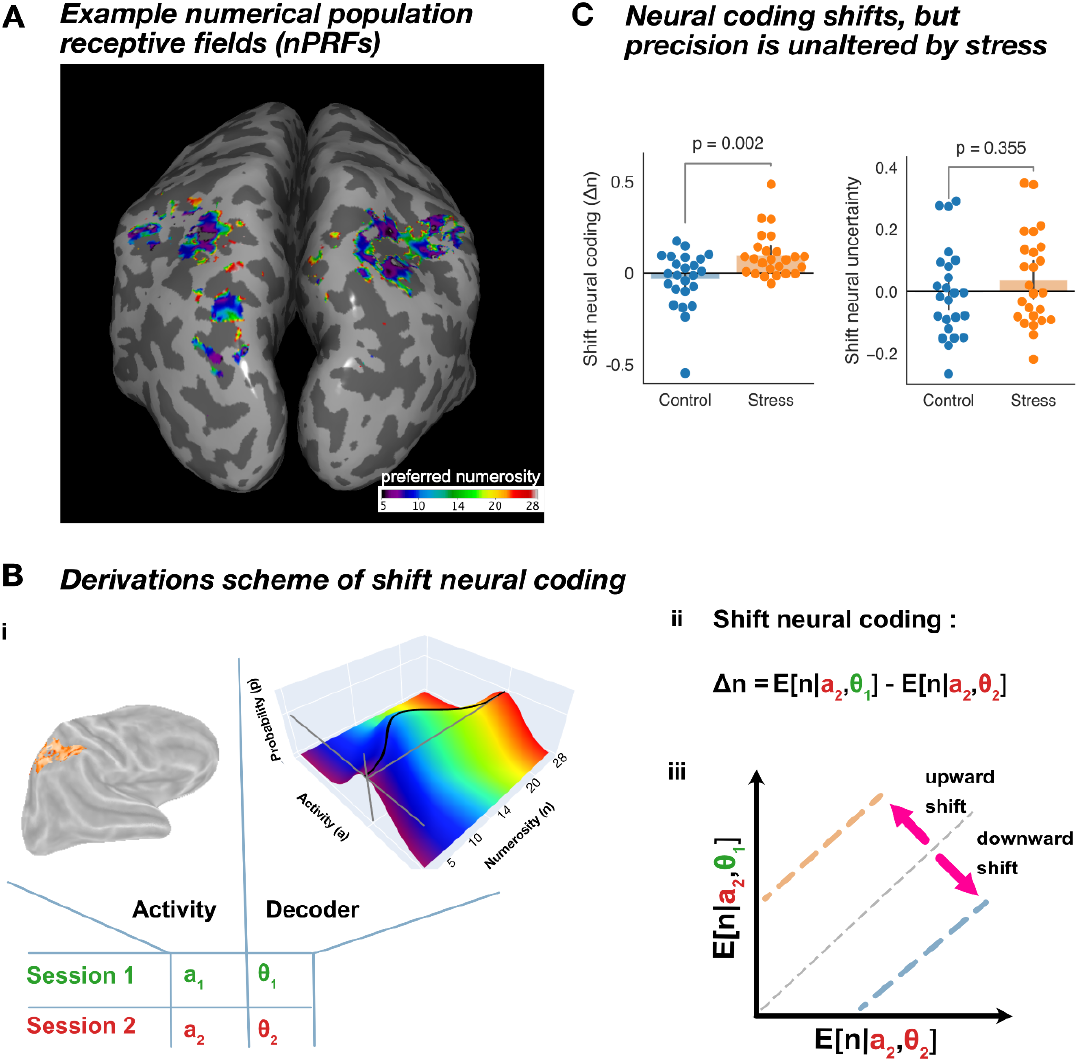
Effect of stress on neural noise and bias. **A. Preferred numerosities of numerical population receptive fields (nPRF) for a representative subject, visualized on an inflated brain surface.** The visualization of RF location was masked by an R2 cutoff (>0.05; see also^33^). Note that most receptive fields lie within the area of the intraparietal sulcus. **B. Quantification of the shift in neural coding across both sessions. i)** nPRF and noise covariance parameters (**θ**) were estimated with data from either session 1 or session 2. These parameters span up a joint probability distribution over multivariate brain activity and payoff magnitudes (the black curve lying on top of the probability surface illustrates the tuning function of that cortical location). Crucially, we can decode a presented numerosity using this joint probability distribution by taking the expectation over the posterior on possible numerosities, given observed brain activity patterns (p(n|a)). These expectations will be different depending on which parameters are used (i.e., those inferred from session 1 or session 2). **ii)** The shift in neural coding (**Δn**) comprises the average difference (over trials) in the decoded numerosity between decoders that were used across (**E[n**|**a**_**2**_,**θ**_**1**_]) versus within (**E[n**|**a**_**2**_,**θ**_**2**_]) sessions. Note that decoders were trained either on session 1 or 2 and always used to decode brain activity during session 2, so that differences between across-session and within-session decoding indicate a shift in the neural code between sessions . **iii)** If the across-session decoder consistently over- (**Δn** >0) or underestimates (**Δn** < 0) the presented numerosity as compared to the within-session decoder, then neural payoff magnitude representations must have shifted from session 1 to session 2. **C. Shift in neural coding and precision of neural magnitude representations.** For the control group, the shifts in neural coding for different subjects are centered around 0. In contrast, for the stress group, they are reliably larger than 0, indicating that a decoder trained on session 1 systematically overestimates presented numerosities when used on neural data of session 2, but only for the stressed group. In contrast, neural uncertainty/decodability - the correlation between the ground truth and the decoded payoff magnitudes - did not shift differently between sessions for the two groups.

We used this fitted model to measure changes in the noisiness of neural number representations and/or shifts in prior beliefs as encoded by the approximate number system. For the former, we correlated the presented numerosity (ground truth) with the ‘neurally decoded’ numerosity over trials for each subject, which has previously been shown to relate to behavioral precision of number representation^33,36,50^. For measuring perception-related prior shifts (as predicted by the change in bias of the cognitive model), we derived a measure that assesses the degree to which neural coding of numbers (i.e., the posterior estimate that combines priors and evidence) shifts in line with the more optimistic perception of magnitudes as evident in our cognitive modelling. This measure was calculated by decoding presented payoff magnitudes from the neural activity in session 2 (during the stress/control task) using two fitted models: The model fitted to the data from session 1 (across-session decoder, in the absence of stress/control, which therefore represents the way the unstressed brain usually interprets the activity patterns in parietal cortex), and the model fitted to the data of session 2 (within-session decoder, the way in which the stressed brain interprets these activity patterns). Any robust differences between these two decoded estimates thus indicate that (and how) the stress/control manipulation during session 2 has shifted neural magnitude coding relative to the unstressed state in session 1. If this difference is positive, the across-session decoder (and thus probably also the brain during stress/control) systematically overestimates the numerosity, implying that the neural code for payoff magnitudes has shifted in line with more optimistic priors.

This pattern is exactly what we found: For the control group, the shifts in neural coding of the individual subjects were centered around zero (t(23)=-1.070, p=0.296, 95%CI=[-0.100, 0.030]), implying that the control manipulation did not systematically affect neural magnitude coding. For the stress group, the shifts were significantly larger than zero (p=0.001, t(24)=3.940, p=0.296, 95%CI=[0.050, 0.150]) and consistently larger than in the control group (t(43.75)=-3.26, 95%CI=[-0.21, -0.05], p=0.002) (Figure 4C). This pattern was evident for both the risky and the safe option (safe: t(45.18)=-3.127, p=0.003, 95%CI=[-0.22 -0.05]; risky:, t(44.63)=-3.122, p=0.003, 95%CI=[-0.21 -0.05]), providing a robustness check and showing that the stress effects applied in general, to the neural coding of both larger uncertain and smaller certain options. In contrast, neural uncertainty (the accuracy of the decoder, related to the precision of number representation) did not change for either group in a specific direction (t(46.97)=-0.934, 95%CI=[-0.12, 0.05], p=0.35). This is again consistent with the findings from the cognitive model, which also indicated no change in the precision of mental payoff representations.

### Prefrontal value signals represent the perceptually shaped magnitude representations

A plethora of neuroeconomics research has revealed that in many economic choice paradigms, the relative subjective value of presented choice options is correlated with neural activity in the medial prefrontal cortex (mPFC) and orbitofrontal cortex (OFC)^67^. Building on this literature, we examined whether neural signals in these regions reflect the perceptually shaped magnitude estimates derived from our Perceptual Memory-based Choice (PMC) model. Specifically, we asked whether the BOLD signal in mPFC and OFC correlated with the magnitude percepts predicted by the PMC, and how these correlations compared to those with more traditional estimates of subjective value, like EV and expected utility..

To examine these correlations, we conducted small-volume corrected analyses within previously proposed subjective-value regions (mPFC from MarsAtlas^71^), as well as whole-brain analyses. We tested for correlations of brain activity with three putative estimates of value of the option (presented at that moment): Objective expected value (EV; objective probability times objective payoff), classical subjective expected utility (EU, as estimated by a standard EU model with a –usually– concave power function^40^), and a *perceived* expected-value measure (pEV) derived from our PMC model. Note that both the classical EU and our new pEV already incorporate any stress-effects, since they were derived from the respective models fit to the choice data in the different conditions. However, only with our new pEV estimate did we find a cluster in the proposed value regions (peak: xyz=[-4,40,17.5], t[DOF]=4.87; cluster size(mm3) = 2025], see Figure S4) that survived statistical thresholding (cluster forming threshold: p<0.001; family-wise error rate at cluster level: α < 0.05). This suggests that prefrontal (subjective) value signals more closely correspond to the cognitive representations embedded in our perceptual model than to EV or EU. Moreover, we only found prefrontal subjective value representations during the presentation of the second option value when participants had to make a choice, not during the first stimulus presentation where they simply encoded the magnitude of that one stimulus for the later comparison. This is consistent with most studies of value-based decision-making, which have reported mPFC subjective value representations at the time of decision-making and thus action selection ^68–70^. Thus, our results suggest that value-coding regions such as OFC/mPFC integrate perceptual-level representations of magnitude into higher-order value signals that guide choices, supporting a link between perceptual encoding and valuation processes.

## Discussion

How stress affects risky financial decision-making has been of considerable interest to various scientific disciplines^10,11^. However, previous studies on the topic have reported contradicting results^22,72^, and a mechanistic understanding of these results is still lacking. To address this gap, we elucidated the underlying neurocognitive processes by which acute stress alters financial decision-making under risk using a novel, *perceptual* account of risk preferences^51^. Using this novel framework, we could disentangle the effects of stress on representational noise from its impact on prior beliefs. This showed that participants became more risk-seeking under acute stress, changing their behavior from slight risk aversion to more risk-neutral choices, but leaving the consistency of choices unchanged. Further computational modeling indicated that this behavioral change was consistent with a model where the participant’s priors became more closely aligned with the objective payout distribution, whereas perceptual and working memory noise were unaffected. Critically, our neural findings are fully consistent with such a shift in prior beliefs, as we observed a concurrent *systematic upward shift in parietal neural coding* of magnitudes under stress, but no difference in neural noise.

Previous studies have reported contradictory findings on the influence of stress on risk preferences, with some suggesting increased risk-seeking^25,72,73^ and others reporting risk aversion^22,74^. A key challenge in interpreting these results lies in the methodological variability across studies, particularly in how risk preferences are measured and conceptualized. Our study takes a different approach by adopting a perceptual perspective, which differentiates between noise and bias in risky financial choice. This account frames risk aversion as a systematic underestimation of the risky option (which always has a larger magnitude than the safe option), as a consequence of Bayesian perceptual inference on the payoffs^30,31,33^. It is precisely this misperception that acute stress appeared to influence in our experiment. As the prior(s) shifted toward larger magnitudes, the perceived distortion of risky options decreased, making them more likely to be chosen.

In other words, by aligning the prior more closely with the actual reward distribution, stress reduced the numerical bias in the risky financial choice. This perceptual perspective on choices about risk allows for a meaningful use of the term ‘adaptive’ -not in a normative sense of what risk attitudes should be, but in the sense that the perceptual encoding of decision-relevant information becomes more veridical under stress (and the associated behavior therefore leads to a higher payoff on the long run). A more veridical prior is consistent with findings that for perceptual decision-making, arousal entails that internal representations become more sensitive to local statistics of the present environment and rely less on long running, global statistics.^75,76^^77^^75,76^

Stress elicits a complex array of physiological and neural responses, encompassing two distinct but parallel systems: the hypothalamic-pituitary-adrenal (HPA) axis and the autonomic nervous system (ANS)^77^. While both systems are activated by stress, they operate on different timescales and have distinct physiological effects. The HPA axis primarily regulates longer-term metabolic and cognitive adaptations to stress, whereas the ANS exerts more immediate influences, with an instantaneous shift in balance between sympathetic and parasympathetic activity^77^. Given that our stress manipulation and behavioral task took place within a single experimental session (approximately one hour), the timescale of our effects is more consistent with modulation by autonomic activity—particularly increased sympathetic tone—than with delayed HPA-axis responses. Consistent with this interpretation, other studies have reported shifts in the perception of magnitudes as a function of sympathetic and parasympathetic tone. Belli et al.^78^ report that both numerosity perception (from dot clouds) and numbers randomly generated by participants tend to be larger during inhalation compared to exhalation. Similarly, Meck and Church^79^ showed that the (sympathetic nervous system) stimulant methamphetamine can shift the psychophysical functions of number and time discrimination leftwards (towards larger values) in rats. Another recent study in humans reported altered time perception under stress, again in a direction of reproducing stimuli longer than before^80^. Thus, it is very well possible that the effects of stress we find are mediated by the autonomic nervous system, and specifically the sympathetic nervous system.

However, stress is a multifaceted phenomenon that can be elicited in different ways and on different time scales, likely involving different physiological systems. The current study investigated only one specific type of stress in a specific type of risky choice paradigm. It is well possible that the effects of acute stress on risky financial decisions we delineate here change substantially in a state of *chronic stress*. For instance, there may be compensatory mechanisms in the perceptual apparatus (that underwent the prior shift under acute stress) that could give rise to opposite effects, more pessimistic priors and thus increased risk aversion, as found in some risk studies examining effects of more long-term stress^10,74^. Future work might clarify this, by applying the perceptual framework we use here in the context of long-term (or non-social types of) stress. Likewise, it is hard to say how the observed effect might translate to other kinds of risky choices, as risk attitudes often do not generalize across different domains^81–83^. Having said that, our findings suggest that stress has very specific effects on numerical perception, by changing neural tuning for magnitudes in the parietal cortex, implying that similar effects are likely to emerge in all types of decisions where the processing of numerical information plays a central role (such as decisions based on information about payoffs time durations, or probabilities).

Beyond the specific context of stress and risky decision-making, our findings also speak to more general questions about how Bayesian perception is neurally implemented—both in the domain of numerosities and beyond. Prior work has shown that the precision with which numbers are neurally encoded in parietal cortex relates to individual differences in numerical acuity^33,36,38,50^, suggesting that behaviorally relevant information is represented in these parietal association areas. However, it has so far remained unclear whether these neural codes reflect raw sensory evidence or posteriors that already incorporate prior beliefs^38^. Our results suggest that prior information does already impinge on relatively ‘early’ numerosity representation in the parietal cortex. Earlier studies that investigated whether the brain represents priors versus posteriors, in other perceptual domains, relied on a *correlational approach*^*84*^, assuming linear relationships between uncertainty and BOLD activity^85–87^. These studies report widespread sensitivity to environmental uncertainty across sensory, frontal, and subcortical regions, but they do not clarify where or how prior information is represented. In contrast, we used a *code-driven framework*^*84*^ based on a probabilistic population code, which, unlike a linear code, can explicitly represent both stimulus likelihood and uncertainty in one and the same neural population^88^. This approach allowed us to disentangle potential shifts in neural noise and prior beliefs, revealing clear evidence for the latter—fully consistent with our cognitive modeling.

Our finding of shifted numerosity coding also reveals interesting parallels with visual attention research, where covert shifts in attention are known to induce shifts in visuospatial receptive field positions^89^. We speculate that numerical receptive fields may similarly shift along the number line toward currently relevant values—an elegant neural mechanism for Bayesian range adaptation. This raises new questions about the origin of such shifts and whether prior and likelihood information map onto distinct cortical depths, in line with canonical laminar patterns^90–92^. To better understand Bayesian coding of numerosity and other abstract features, future studies should manipulate both priors and stimulus reliability. Combining numerical population receptive field mapping with a code-driven analysis could reveal how prior beliefs and sensory evidence are represented across the brain. Multiple cortical areas encode numerosity with non-linear tuning; it remains to be tested whether these regions differ in their sensitivity to prior information. Interestingly, visuospatial tuning shifts are typically coordinated across areas to maintain stable reference frames^93,94^. Whether similar coordination exists in higher-order domains like number remains an open and intriguing question.

A limitation of our study is that, because a large majority of participants in the stressed group (21 out of 25) were risk-averse or risk-neutral during the first session, we cannot effectively distinguish between two different but highly related hypotheses: (a) stress induces an upward shift in prior beliefs over payoffs, or (b) stress leads to more veridical priors that better reflect the actual payoff distribution. These two hypotheses may be disentangled by future studies that include more participants with risk-seeking preferences, or that explicitly manipulate payoff priors to assess how environmental statistics interact with the stress manipulation. Our finding that shifts in bodily state, as induced by stress, have a profound influence on perception and thereby (economic) decision-making may have clinical implications as well. For example, stress is a known perpetuating factor in excessive gambling behaviors, and our findings may offer a perceptual explanation for this link:. Stress may lead to an overly optimistic (mis)perception of risky gambles, thereby affecting perceived payoff magnitudes and/or probabilities. This aligns with evidence that problem gamblers tend to be overconfident in risky bets^95^. Thus, further exploring the interplay between stress, numerical perception, and gambling behavior could refine models of decision-making mechanisms underlying problem gambling. Notably, such a theoretical shift from valuation to perceptual processes might also lead clinical research to employ different experimental paradigms and focus on pathological changes in areas outside of the classical valuation network (e.g., the parietal lobe). Similar translational opportunities exist in anxiety disorders, where chronic stress is a common trigger. Patients with anxiety often exhibit heightened sensitivity to losses and uncertainty, along with more pessimistic expectations about potential adverse events and outcomes^96^. A perceptual, Bayesian lens on these two behavioral markers might mechanistically explain how perceptual distortions contribute to maladaptive decision-making in anxiety disorders, over and above affective and/or valuation.

### Conclusion

To conclude, our study can serve as a stepping stone to better understand the neuro-cognitive processes that underlie stress-related changes in risk attitudes. Our cognitively interpretable psychophysical and neural models may provide handles to reconcile opposing findings of earlier stress-risk studies, and may contribute to helping people make better choices in challenging situations that comprise uncertainty.

## Methods

### Participants

Fifty healthy male participants (18 to 30 years old) underwent the complete scanning procedure. We invited only male participants to reduce the variance of the hormonal stress response, as it might be altered in females by the menstrual cycle and its associated fluctuations in gonadal hormones^97^. Additional exclusion criteria were prescription medication taken within the last 2 months, smoking regularly (at least 1 cigarette per day), or the regular use of illegal substances. The first session was initially performed with 59 participants, hence 9 participants did not complete the second scan for several reasons (6 participants voluntarily withdrew from the study, including 1 subject during the second scan, and three others were excluded for not meeting the eligibility criteria for the second session). We also screened participants for MR compatibility prior to their participation in the study. To ensure unbiased choices and stress responses, we neither recruited participants who studied psychology or economics, nor those who had taken part in previous stress-induction studies. All participants were right-handed, had no indications of psychiatric or neurological disorders, did not need visual correction, and had normal color vision. They were instructed to not drink alcohol, smoke, take drugs or medication, and abstain from vigorous physical 24 hours before the study. Moreover, they had to abstain from food, caffeine, chewing gum, and any drinks other than water for 2 hours before the experiment, and had to have had enough sleep (same duration as their habitual sleep and a minimum of 6 hours) in the night before the study. Session 2 scanning was restricted to 12 - 6 pm due to the daily cortisol cycle. Our experiments conformed to the Declaration of Helsinki and our protocol had the approval from the Canton of Zurich’s Ethics Committee.

### Procedure

Participants came to the laboratory twice, once for the baseline session and once for the experimental stress/control session. During the first session, they received a written instruction about the experiment. Next, they went into the scanner and performed a calibration version of the risky choice task while anatomical scans were acquired. Participants had to make a decision about 96 choice problems, always choosing between either a safe amount of money or a substantially larger potential payoff with a 55% chance of payout (and 45% chance of winning nothing). The safe payoff was either 7, 10, 14, 20, or 28 (Swiss francs), while the risky option was 2^*h*/4^ times the safe option with *h* being all integers between 1 and 8. These gambles were presented twice, alternating the presentation order of the safe and risky option. Utilizing a probit model, we analyzed the choice data with “choosing the risky option” as the dependent variable and an intercept alongside the log-ratio (risky/safe option) as the independent variables (see also ^30,33^). The model effectively related risky-to-safe payoff ratios with the likelihood of selecting the risky option. Subsequently, a personalized design was created for each participant, consisting of 6 logarithmically spaced fractions which, according to the calibration data fits, should lead to risky choice proportions ranging from 20% to 80%, enabling optimal data for more complex behavioral modeling for later on. These fractions were combined with 5 safe payoff levels (7, 10, 14, 20, 28) and presented four times in each session, alternating the order of presentation between safe and risky options to ensure equal exposure to all permutations. This resulted in a total of 120 trials per session that were split up into 6 runs, lasting about 6 minutes each.

### Risky choice task paradigm

The subject’s task was to select between a certain amount of money (7/10/14/20 or 28 CHF) or a gamble with a 55% probability of winning a larger amount of money and a 45% probability of winning nothing at all. These choices were presented through a series of customized stimuli, as depicted in Figure 1C. Each screen displayed a red cross with two diagonal lines to maintain fixation near the center and prevent any confusion regarding the numerosity of stimuli that a standard fixation cross or point might cause. The start of each trial was signaled by the fixation cross turning green for 250 ms. Following a 300 ms pause with only the red fixation cross, a pie chart with a diameter of 1 degree-of-visual-angle (dova) indicated the probability of payout for the forthcoming stimulus, which was consistently either 55% or 100%. After an additional 500 ms of fixation, a display of 1-CHF coins representing the potential payoff of the initial choice option was presented for 600 ms. These coins, with a radius of 0.3 dova, were randomly positioned within a circular aperture of 5.25 dova. Afterward, only the fixation cross was displayed for a variable duration ranging from 5 to 8 seconds. Following this, another pie chart indicating the probability of the second payoff was shown for 300 ms, succeeded by a 300 ms fixation screen, and then another display of coin stimuli representing the potential payoff of the second option, again for 600 ms. Subjects had to indicate their response using their index finger for the first-presented option or their middle finger for the second-presented option. Upon responding, they were presented with a 1 or 2, at the center of the screen, for 500 ms to give them feedback about their choice. After the second coin stimulus, there was an inter-trial interval of 4, 4.5, 5, or 5.5 seconds, showing only the fixation cross. Stimuli were presented using a projector on a white screen at the back of the bore and a hot mirror system on top of the coil system (1920 x 1080 pixels). The projector screen was at a distance of 125cm from the subject’s eyes and was 42 cm wide, corresponding to a field-of-view of approximately 19 DoVA.

For incentive compatibility, after participants finished all trials, one trial was randomly drawn for the payout. If on that trial a participant had chosen the certain option, she immediately received that amount. If she had chosen the risky option, she had to roll a virtual 100-sided die. Any result smaller than or equal to 55 was paid out, any other result led to no payout.

### Stress and control calculation task

For the second session, subjects randomly assigned to the stress group underwent an adapted version of the Montreal Imaging Stress Test (MIST, ^60,64,98^) for which they had to solve challenging mental arithmetic questions under time pressure. Specifically, we employed an adaptive task that gradually increased in difficulty when participants achieved a correct response rate above 50% and decreased in difficulty when their performance fell below this threshold. In the stress group, after each trial participants got feedback on whether their answers were correct and additionally, they received visual (false) feedback about their performance relative to the group average of all other study participants prior to their session. Participants were informed that maintaining a similar level of cognitive effort was crucial for their data to be usable. They were instructed to keep their performance within a green zone on a performance bar and avoid dropping into a red zone, as shown in Figure 1D. To further increase stress levels, we periodically interrupted the scanning to remind participants that they needed to improve their performance to ensure their data could be used for further analysis. Additionally, participants were led to believe that experimenters were qualitatively assessing their performance through a bidirectional video stream. Crucially, experimenters provided social evaluative feedback on the screen through critical facial expressions while participants performed the task. (see Figure 1D). Subjects in the control group had to solve the same mental arithmetic questions from the MIST but without the deceptive feedback, social evaluation and the time pressure. Three 6 minutes of the MIST/control task blocks then interleaved the risky choice task runs (see Figure 1B). Lastly, subjects in the experimental group were debriefed.

### Stress measures

To assess cortisol responses, saliva samples were gathered throughout the experiment using oral swabs (Sarstedt Salivette), which participants held in their mouths for about 2 minutes while gently chewing on them. A total of seven saliva samples were collected, with timing relative to the onset of stress induction: Upon arrival (sample 1; –40 minutes), before the first MIST (sample 2; 0 minutes), during the second MIST (sample 3; ∼20 minutes), during the third MIST (sample 4; ∼ 37minutes), at the conclusion of the scan (sample 5; 51 minutes), and following the debriefing session (sample 6; 65 minutes). Post-collection, saliva samples were frozen at –20°C until analysis. Participants were also asked to rate their perceived stress levels using a visual analogue scale during each saliva collection, ranging from ‘not at all’ (1) to ‘very much’ (7). Additional stress ratings were obtained after the first MIST (rating 3; 5 minutes), but saliva samples were not taken at these intervals (see Figure 1D). Upon thawing, the oral swabs were centrifuged at 3000 rpm for 5 minutes, yielding a clear supernatant with low viscosity. Salivary cortisol concentrations were determined using a commercially available chemiluminescence immunoassay with high sensitivity (IBL International, Hamburg, Germany). Both the intra- and inter-assay coefficients for cortisol were below 9%.

The salivary cortisol concentrations over the timepoints were analysed with the area under the curve (AUC) approach^64,99^ to obtain one cortisol measure per participant. Samples from one participant were not usable by the analyzing lab due to the low amount of saliva provided. Consequently, the final analysis included 49 participants (24 in the stress group and 25 in the control group) whose saliva samples were analyzed.

Additionally, we analyzed heart rate and heart rate variability (HRV) measures that have been related to autonomic nervous system (ANS) functioning^61,62^. We analyzed the electrocardiogram (ECG) data that was also collected during fMRI scans with the NeuroKit2 toolbox, which provides processing routines for neurophysiological signals^100^. We used the *v1* column of the physio.log file provided by the MR Scanner’s ECG device and applied the processing steps as recommended by the toolbox (*ecg_process, ecg_peaks, hrv_frequency*), which filters and detrends the ECG signal, finds R-peaks and computes key measures of HRV in both the time and frequency domains (see^100^ for more details on how these measures are computed). We report between-group tests for individual sessions and session differences on subject-wise means of heart rate, standard Deviation of Normal-to-Normal intervals (SDNN - the classical HRV measure), Root Mean Square of Successive Differences (RMSSD) and low frequency high frequency HRV ratio in Figure S2. Before the between-group tests, we tested if values were normally distributed (with *scipy*.*stats*.*normaltest*) and applied either a t-test, which assumes normal distributions, or the non parametric Mann-Whitney U test (for not-normally distributed data).

### Behavioural analyses

#### Bayesian analysis

Our behavioral modeling analyses were done within a Bayesian framework. We used Python packages Bambi^101^ for psychophysical modeling and Bauer^102^ for our mechanistic modeling approach. Both packages are based on pymc^103^, which uses automatically tuned Hamiltonian Markov Chain Monte Carlo sampling to estimate posterior parameter distributions using the No-U-Turn sampling algorithm (NUTS)^104^. We performed formal model comparison using expected log-predictive density (ELPD) measures^66^. When testing differences between parameters, we used Bayesian p-values^105^. Specifically, we took samples from a posterior parameter distribution, or the difference of two parameter distributions and tested which proportion of samples was smaller/larger than 0.

#### Psychophysical model

##### Model specification

As a first behavioral analysis, we used a standard psychophysical (probit) model to dissociate differences in choice consistency (slope) from the indifference point (intercept) ^30,106^. Thus, we fitted a model where the probability of choosing the risky option was a function of an intercept δ and a slope γ times the log-ratio of the risky and safe payoffs:

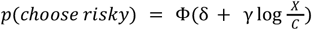

where *X* is the payoff magnitude for the risky option and *C* is the payoff magnitude of the safe option.

##### Parameter estimation

We fitted the psychophysical model using the Bayesian hierarchical GLM-estimation package Bambi^107^. As we were interested in any effect stress might have on the indifference point and choice consistency, we added the effects of session, group and their interaction. Additionally, we also estimated subject-wise random effects on all parameters to obtain subject-wise psychophysical parameters. The following formula was given to the Bambi model estimation procedure:

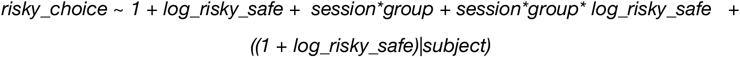

Note that the point-of-indifference of the probit model is a function of both the intercept and slope parameters (-intercept/slope). To meaningfully quantify the point-of-indifference, we used the risk-neutral-probability (RNP) ^37,RNP; 108^. This is the hypothetical probability of the risky option for which a risk-neutral decision-maker would show the same point-of-indifference as the subject. Thus, any RNP above 55% implies risk-seeking, whereas any RNP below 55% implies risk aversion. We defined subjects as being risk averse when the 95% Credible Interval of their RNP was below .55, and risk-seeking if the CI was above .55. When the CI overlapped with .55, we categorized subjects as risk-neutral. The risk neutral probability was derived from the condition- and subject specific slope γ and intercept δ as follows: 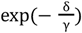.

To fit the hierarchical probit/KLW model, we used the standard weakly informative priors as implemented by default in Bambi (based on the approach of the well-known *rstanarm*-package) package^107^. Briefly, the priors are Gaussian distributions centered at 0, with as a standard deviation 2.5 times the ratio between the standard deviation of the dependent and independent variable ^107^. To sample from the parameter posteriors, we used the no U-Turn sampler (NUTS), which is a self-tuning version of the Hamiltonian MCMC sampler and is particularly efficient for high-dimensional models with correlations between the parameters, such as our model here ^109^. We collected 4 chains of 2000 samples each (1000 burnin). We always visually inspected the traces of the NUTS sampler and made sure the Gelman-Rubin statistic was below 1.05 for all parameters.

#### PMC model

##### Model specification

Our Perceptual Memory-based Choice model (PMC) builds upon an earlier model of Khaw, Li and Woodford ^30,31^ and assumes that subjects base their choices not on the actual payoffs *X* and *C*, but on noisy neurocognitive representations of those payoffs *r*_*x*_ and *r*_*c*_, which are defined as random variables conditional on the actual stimulus values. In the PMRC model, the noisiness of the representation may also depend on the order of presentation:

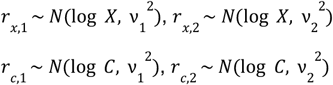

where *r*_*x*,1_ pertains to the representation of the payoff of the risky option presented first, *r*_*x*,2_ to the representation of the payoff of a risky option presented second, *r*_*c*,1_ to the representation of the payoff of a safe option presented first, and *r*_*c*,2_ to the payoff of a safe option presented second. Note that the noisiness of the risky and safe options *ν*_*i*_ is the same, only the order of presentation influences noisiness. We expected that the noise for the first option *ν*_1_ will be higher than that of the second option *ν*_2_, but this is not enforced anywhere in the model (e.g., they have identical priors in the estimation procedure).

The PMRC model also assumes that subjects (potentially) employ different priors for the risky and safe payoffs, because these payoffs come from objectively different distributions:

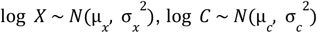

For a given parameter set [*ν* _1_, *ν* _2_, μ _*x*_, μ _*c*_, σ _*x*_, σ _*c*_] and objective payoffs *X* and *C*, we can now obtain the distribution of the expectations of the subject for the two options, which is the product of the prior and evidence distributions, and follows a normal distribution (in logarithmic space) :

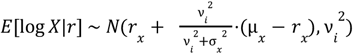

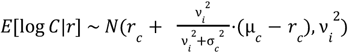

Thus, the distribution of differences between these expectations in log space is given by the difference of these two distributions, *E*[log *X*|*r*] - *E*[log *C*|*r*], which is also a normal distribution:

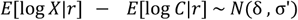

with

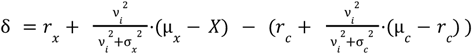

and

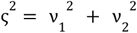

The likelihood of choosing the risky option according to the the PMRC model is defined as the probability that the subject’s estimate of the difference between the risky and safe option (*E*[log *X*|*r*] − *E*[log *C*|*r*]) is larger than the payout probability of the risky option in log space:

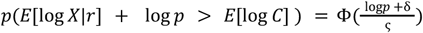

where Φ(*x*) is the standard cumulative normal distribution.

##### Model estimation

We implemented the PMCM using the Bayesian statistical modeling library pymc^v 4.3.0;,110^ and wrapped in our Python package *bauer*^*1*^. We used a hierarchical (regression) approach for estimation. This means that for all main six parameters of the model [*ν* _1_, *ν* _2_, μ _*x*_, μ _*c*_, σ _*x*_, σ_*c*_], we assume a group distribution which is a Gaussian distribution. Furthermore, for the *ν*- and σ -parameters, which are necessarily non-negative, we estimated transformed parameters in an unrestricted space [− ∞, ∞] which we then transformed into the non-negative domain [0, ∞] using the softplus function log(1 + e*x*p(*x*)).

For a given parameter θ (e.g., *ν*_1_ or σ_*x*_), the subject-session-specific parameter of subject *p* at session *t* was a linear combination of the mean group parameter μ_θ_ plus an individually-determined offset-parameter δ_θ_ times the standard deviation of the group distribution σ_θ_ :

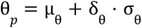

This offset-based specification of the hierarchical model was chosen to prevent the likelihood “funnels” that plague high-dimensional hierarchical models ^111^.

To include potential effects of session and being in the stress manipulation group, we used a regression approach^112^, where a given parameter θ_*p*_ (e.g., *ν*_1_ or σ_*x*_) of subject *p* on a trial *t* (and manipulation group *m*) was a linear sum of both an intercept θ_0_ and multiple regression coefficients θ_*n*_ times their dummy variable *x*, indicating the session and manipulation group assignment (0 or 1):

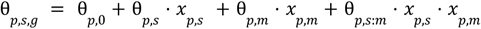

Hence, θ_*p,s*:*m*_ captures any effect related to the stress manipulation (interaction of session and manipulation group). Furthermore, we specified that if any prior mean shifts, both (risky μ_*x*_ and safe μ_*c*_) should shift equally; hence, there was only a common regressor for the means of the priors^2^.

We used mildly informative priors on all group distribution parameters:

For the evidence parameters:

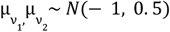 (before softplus transformation)

For the mean of the prior on risky payoffs:

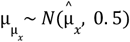

where 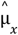 is the *objective* mean of the payoffs of risky options (in log space)

For the mean of the prior on safe payoffs:

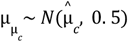

where 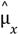 is the *objective* mean of the payoffs of safe options (in log space)

For the standard deviation of the priors:

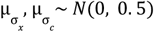 (before softplus transformation)

The prior on all group variance parameters was a Half-Cauchy distribution ^113^:

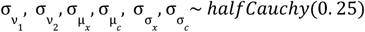

Posterior estimates were obtained using the NUTS-sampler as implemented in pymc 4 using a target acceptance rate of 90%. We obtained 4 chains with 3000 samples each (1500 burnin). If we encountered divergences, we increased the target acceptance rate to 92.5%.

To obtain estimates of the size and presence of an effect, we used “Bayesian” p-values –the probability mass of the posterior estimate above/below 0 ^114^. We generally used one-tailed cutoffs of 5% probability mass.

Model comparisons were performed using the estimated log pointwise predictive density(ELPD)^66^ which is a state-of-the art model comparison technique that – unlike other information theoretic measures like BIC and DIC – takes into account the shape of the posterior distribution and the effective number of parameters of the model.

##### Expected Utility model

We also fitted an Expected Utility model as a baseline model to compare the PMC model against. The EU model assumes that the utility of a payoff is a power function of the payoff:

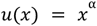

The likelihood of choosing the risky option for a given risky payoff *X* and safe option *C* is then:

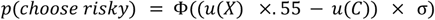

Where Φ is the standard cumulative normal distribution. Just as for the PMC model, this probability is plugged into a Bernoulli likelihood function. α and σ are free parameters, which are fitted using hierarchical Bayesian estimation, using the same *bauer* package as the PMC model was implemented in. We always allowed for a random *session* and *group* effect on both parameters. We then had one version that allowed for an effect of stress (i.e., *group x session* interaction), and one without it.

### MRI data

#### Acquisition parameters

We acquired functional MRI data using a Philips Achieva 3T whole-body MR scanner equipped with a 32-channel MR head coil. For both sessions,we collected 6 runs of fMRI data with a T2*-weighted gradient-recalled echo-planar imaging (GR-EPI) sequence (150 volumes + 5 dummies; flip angle 90 degrees; TR = 2286 ms, TE = 30ms; matrix size 96 × 70, FOV 240 × 175mm; in-plane resolution of 2.5 mm; 39 slices with thickness of 2.5 mm and a slice gap of 0.5mm; SENSE acceleration in phase-encoding direction (left-right) with factor 1.5; time-of-acquisition 4:52 minutes). Additionally, we acquired a high-resolution T1-weighted 3D MPRAGE image (FOV: 256 × 256 × 170 mm; resolution 1 mm isotropic; *TI* = 2800 ms; 256 shots, flip angle 8 degrees; *TR* = 8.0 ms; *TE* = 3.7 ms; SENSE acceleration in left-right direction 2; time-of-acquisition 5:35 minutes).

#### fMRI preprocessing

Preprocessing on fMRI data was performed using fMRIPrep 20.2.2^115^, which is based on Nipype 1.6.1 ^116,117^.

##### Anatomical data preprocessing

The T1-weighted (T1w) images were corrected for intensity non-uniformity (INU) with N4BiasFieldCorrection ^118^, distributed with ANTs 2.3.3 ^119^. The T1w-reference was then skull-stripped with a Nipype implementation of the antsBrainExtraction.sh workflow (from ANTs), using OASIS30ANTs as target template. Brain tissue segmentation of cerebrospinal fluid (CSF), white-matter (WM) and gray-matter (GM) was performed on the brain-extracted T1w using fast ^FSL 5.0.9;,120^. A T1w-reference map was computed after registration of 3 T1w images (after INU-correction) using mri_robust_template ^FreeSurfer 6.0.1 121^. Brain surfaces were reconstructed using recon-all^FreeSurfer 6.0.1;,122^, and the brain mask estimated previously was refined with a custom variation of the method to reconcile ANTs-derived and FreeSurfer-derived segmentations of the cortical gray-matter of Mindboggle ^123^. Volume-based spatial normalization to one standard space (MNI152NLin2009cAsym) was performed through nonlinear registration with antsRegistration (ANTs 2.3.3), using brain-extracted versions of both T1w reference and the T1w template. The following template was selected for spatial normalization: ICBM 152 Nonlinear Asymmetrical template version 2009c^124^,

##### Functional data preprocessing

For each of the 12 BOLD runs found per subject (across both sessions), the following preprocessing was performed. First, a reference volume and its skull-stripped version were generated using a custom methodology of fMRIPrep. BOLD runs were slice-time corrected using 3dTshift from AFNI 20160207^125^. Head-motion parameters with respect to the BOLD reference (transformation matrices, and six corresponding rotation and translation parameters) are estimated before any spatiotemporal filtering using mcflirt FSL 5.0.9^FSL 5.0.9;,126^. A B0-nonuniformity map (or fieldmap) was estimated based on two (or more) echo-planar imaging (EPI) references with opposing phase-encoding directions, with 3dQwarp^125^ (AFNI 20160207). Based on the estimated susceptibility distortion, a corrected EPI (echo-planar imaging) reference was calculated for a more accurate co-registration with the anatomical reference. The BOLD reference was then co-registered to the T1w reference using bbregister (FreeSurfer) which implements boundary-based registration^127^. Co-registration was configured with six degrees of freedom. The BOLD time-series (including slice-timing correction when applied) were resampled onto their original, native space by applying a single, composite transform to correct for head-motion and susceptibility distortions. These resampled BOLD time-series will be referred to as preprocessed BOLD in original space, or just preprocessed BOLD. The BOLD time-series were resampled onto the following surfaces (FreeSurfer reconstruction nomenclature): fsaverage, fsnative. Several confounding time-series were calculated based on the preprocessed BOLD: framewise displacement (FD), DVARS and three region-wise global signals. FD was computed using two formulations following Power absolute sum of relative motions^absolute sum of relative motions, 128^ and Jenkinson (relative root mean square displacement between affines ^126^. FD and DVARS are calculated for each functional run, both using their implementations in Nipype (following the definitions by Power et al. 2014). The three global signals are extracted within the CSF, the WM, and the whole-brain masks. Additionally, a set of physiological regressors were extracted to allow for component-based noise correction CompCor^CompCor, 129^. Principal components are estimated after high-pass filtering the preprocessed BOLD time-series (using a discrete cosine filter with 128s cut-off) for the two CompCor variants: temporal (tCompCor) and anatomical (aCompCor). tCompCor components are then calculated from the top 2% variable voxels within the brain mask. For aCompCor, three probabilistic masks (CSF, WM and combined CSF+WM) are generated in anatomical space. The implementation differs from that of Behzadi et al. in that instead of eroding the masks by 2 pixels on BOLD space, the aCompCor masks are subtracted from a mask of pixels that likely contain a volume fraction of GM. This mask is obtained by dilating a GM mask extracted from the FreeSurfer’s aseg segmentation, and it ensures components are not extracted from voxels containing a minimal fraction of GM. Finally, these masks are resampled into BOLD space and binarized by thresholding at 0.99 (as in the original implementation). Components are also calculated separately within the WM and CSF masks. For each CompCor decomposition, the k components with the largest singular values are retained, such that the retained components’ time series are sufficient to explain 50 percent of variance across the nuisance mask (CSF, WM, combined, or temporal). The remaining components are dropped from consideration. The head-motion estimates calculated in the correction step were also placed within the corresponding confounds file. The confound time series derived from head motion estimates and global signals were expanded with the inclusion of temporal derivatives and quadratic terms for each^130^. Frames that exceeded a threshold of 0.5 mm FD or 1.5 standardised VARS were annotated as motion outliers. The BOLD time-series were resampled into standard space, generating a preprocessed BOLD run in MNI152NLin2009cAsym space. First, a reference volume and its skull-stripped version were generated using a custom methodology of fMRIPrep. All resamplings can be performed with a single interpolation step by composing all the pertinent transformations (i.e. head-motion transform matrices, susceptibility distortion correction when available, and co-registrations to anatomical and output spaces). Gridded (volumetric) resamplings were performed using antsApplyTransforms (ANTs), configured with Lanczos interpolation to minimize the smoothing effects of other kernels^131^. Non-gridded (surface) resamplings were performed using mri_vol2surf (FreeSurfer). First, a reference volume and its skull-stripped version were generated using a custom methodology of fMRIPrep. BOLD runs were slice-time corrected using 3dTshift from AFNI 20160207^125^. The BOLD reference was then co-registered to the T1w reference using bbregister (FreeSurfer) which implements boundary-based registration^127^. Co-registration was configured with six degrees of freedom. The BOLD time-series (including slice-timing correction when applied) were resampled onto their original, native space by applying the transforms to correct for head-motion. These resampled BOLD time-series will be referred to as preprocessed BOLD in original space, or just preprocessed BOLD. The BOLD time-series were resampled onto the following surfaces (FreeSurfer reconstruction nomenclature): fsaverage, fsnative. The BOLD time-series were also resampled into standard space, generating a preprocessed BOLD run in MNI152NLin2009cAsym space. Many internal operations of fMRIPrep use Nilearn 0.6.2^132^, mostly within the functional processing workflow. For more details of the pipeline, see the section corresponding to workflows in fMRIPrep’s documentation.

#### fMRI analysis - multivariate

The main goal of our fMRI analyses was to get a neural surrogate of the noise and the bias (prior) with which numbers are processed in the brain and how that might be altered by stress.Following earlier work^108,133,134^, we used an encoding/decoding-modelling approach, where we inverted an encoding model that describes how a voxel *i* responds to specific stimulus magnitude *s f*(*s*)→*x_i_* in a Bayesian framework, by extending the encoding model to a multivariate likelihood function *p*(*X*|*s*) using a multivariate t-distribution: [*x*_1_, .., *x*_*n*_] ∼ f_1..n_(*s*) + ϵ with ϵ ∼ *t*(0, Σ, *d*), where Σ is the residual covariance and *d* is the degrees-of-freedom of the t-distribution. Using an explicit likelihood function allowed us to decode from trial to trial what was the presented payoff magnitude, as well as the fidelity of the neural response. In essence, the model allows us to, for every trial, assess the consistency between the observed BOLD activation pattern and the hypothesized neural representations of numerosity. The fidelity of the neural response was operationalised as the dispersion (standard deviation) of the decoded posterior. Intuitively, the posterior would be less dispersed when the BOLD responses of many voxels agree with a small set of numerosities. When the amplitude of the response across voxels is smaller and/or they are not consistent with the same numerosities, the posterior will be more dispersed, as the decoder is less certain about which stimulus was presented. We postulate, as others have done before ^133,134^, that the uncertainty of our decoding of the presented stimulus is related to the fidelity of the *neural representation* of that stimulus.

The main fMRI analysis can be split up in the following steps: 1) Fit a single-trial GLM to estimate trialwise measures of the response amplitude across voxels 2) Fit a numerical receptive field model ^135^ to the response, 3) Fit a multivariate noise model to the residuals of the nPRF model in a leave-one-run-out cross-validation scheme ^133^, 4) Obtain a posterior estimate of the payoffs magnitudes of unseen data using the noise model and an inverted nPRF model, 5) Derive the measure of neural noise by correlating the expected value of the posterior estimates with the true numerosities of the respective trial 6) Derive the measure of shift neural coding from the average difference of the across versus within session decoding estimates.

##### Single trial estimates

We used the GLMSingle Python package ^136^ to obtain single-trial BOLD estimates. Briefly, the GLMSingle package uses cross-validation to do model selection over GLMs with a) a library of different hemodynamic response functions b) different L2-regularisation parameters to shrink the single trial estimates, combating the issue of correlated single trial regressors ^137^. c) GLMSingle also obtains GLMDenoise ^138^ regressors based on the first *n* PCA components in a set of noise voxels. Noise voxels are defined as having low explained variance in the task-based GLM. The number of PCA components is selected via cross-validation.

As input to GLMSingle, we used both the 24 first and 24 second payoff presentations per run. For the second payoff presentations, we modeled all trials with the same numerosity as being in the same condition, to aid GLMSingle with cross-validation (similar numerosities should have similar responses). Because stress manipulations can have strong physiological consequences (e.g. increased heart rate), we decided to include additional confound regressors generated with the RETROICOR method^139^, implemented in the matlab based TAPAS software collection^139,140^, accounting for effects of respiration and cardiac pulsatility on fMRI signal.

##### Numerical receptive field modeling

As a first step, we fitted a numerical receptive field model to all voxels in the brain, for each session separately (so using 120 single trial estimates). Although this method is described in detail elsewhere^108,135^, we briefly describe it here. First, we estimate the mean μ, standard deviation σ, amplitude *A* and baseline *B* of a log-normal receptive field for every voxel in the brain separately, to predict the BOLD response of these voxels to different numerosities.

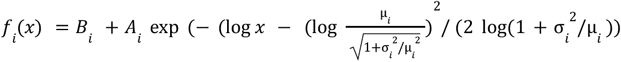

The second part of this equation is a parameterisation of the log-normal probability density function where μ and σ are the mean and standard deviation of the distribution in natural space. Although this parameterisation is somewhat exotic, it is highly useful when plotting the estimated preferred numerosities and their dispersion, for example on the cortical surface.

We fitted the model by first correlating the single trial estimates for the first payoff stimulus presentation with the predictions of a large grid of 60 μ-s between 5 and 80 and 60 σ-s between 5 and 40. We then estimated *I* and *A* using linear least-squares on the best-correlating μ and σ-parameters. Finally, we used gradient descent^141^ to refine parameters further.

##### Voxel selection

We used a nested cross-validation scheme to select the voxels to decode from for each subject/run similar as described in de Hollander 2024^36^, which avoided the arbitrary choice of the number of voxels within the IPS area. Voxel selection was restricted to individual right numerical parietal cortex (rNPC) masks, obtained by warping a group-average rNPC mask (as in n^33^), based on average *R*^2^ -maps from the nPRF model described above) from *fsaverage*-surface space to individual surface and then volume (anatomical) space using *neuropythy* (https://github.com/noahbenson/neuropythy).

Specifically, when decoding run *i* (out of a total of 6 runs per session), we fitted the nPRF model on all voxels within the NPC mask on (4-run) subsets of the remaining 5 runs, by leaving another run *j* out *R*_i*j*_. Then, for each of these nested folds *R*_i*j*_, we calculated the *out-of-sample R*^2^ on run *j* for each voxel. Thus, for each voxel and each decoded test run *i*, we had 4 out-of-sample *R*^2^ *s*, which we averaged over. For decoding run *i*, we only used voxels that had a mean out-of-sample *R*^2^ larger than 0.

##### Decoding

After voxel selection, we used a leave-one-run-out cross validation scheme where the nPRF model was fitted to all runs but the test run, after which also a multivariate noise model was fitted to the residual signals. Specifically, we fitted the following covariance matrix:

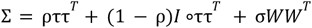

as well as the degrees-of-freedom of a 0-centered, multivariate t-distribution. Briefly, τ is a vector as its length the number of voxels (*n*) in the ROI. It pertains to the standard deviation of the residuals of each voxel. Thus, ρ determines to which extent all voxels correlate with each other (ττ^*T*^ is the covariance matrix of perfectly correlated voxels, whereas *I* °ττ^*T*^ corresponds to a perfectly diagonal matrix/spherical covariance). *W* is a square matrix of *n* ×n. Each element *W*_*i,j*_ is the product of the receptive fields of voxels *i* and *j* across stimulus space (*f*_*i*_ (*S*) *f*_*j*_ (*S*)^*T*^ with *S* being the entire stimulus space [5, 6, .., 112]. Thus, the free scalar parameter σ determines to which extent voxels with overlapping receptive fields have more correlated noise.

Once the noise model was fitted, we set up a likelihood function for any multivariate BOLD pattern *X* and for any numerical stimulus *s*:

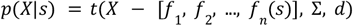

Since we assumed a flat prior on all integers between 5 and 112, we evaluated this likelihood on all these integers and normalized the resulting discretized probability density function (pdf) to integrate to 1. We took the expected value of this pdf to be our estimate of the presented stimulus. Decoding accuracies of individual subjects were then obtained by taking the coefficient of the correlation between those estimates with the true numerosity presented at the respective trials, thus, theoretically ranging from -1 to 1. This was our measure of neural precision, as previous studies have shown that it relates to behavioral precision from numerical tasks^50,33^

##### Neural stress effects: Shift in neural coding

Apart from quantifying neural noise (as the inverse of neural precision), we also examined how stress changed the neural coding of prior expectations. Assuming that activity in parietal nPRFs is a neural representation of the mental percept of a number in which prior expectations have already been integrated (hence, not representing “raw” incoming sensory evidence), we reasoned that any change in prior should result in a corresponding change of parietal number encoding. To test this, we calculated the difference of within versus between-session decoding estimates (expected value of the pdf) for each trial of each subject. In other words, we applied either the (encoding) model parameters from session 1 or session 2 to the betas/activity data (single-trial BOLD estimates) from session 2, to obtain a numerosity estimate of that trial. If neural numerosity coding shifted systematically towards larger or smaller numbers, this shift in neural coding would thus be consistently expressed as a positive or negative misclassification averaged over all trials of that subject. Moreover, in the across-as opposed to the within-session decoding approach, we did not use an leave-one-run-out cross validation scheme, as here all runs of the test datasets were independent from the training datasets. Hence, voxels were selected based on (across all runs averaged) cross-validated *R*^2^ maps and nPRF parameters fitted to all runs from session 1 in order to decode numerosities of session 2 trials.

#### fMRI analysis - univariate

Finally, we wanted to test alternative explanations for the change in behavior, for instance that stress-induced increased risk-seeking would reflect altered subjective valuation and not perceptual processes. Hence, we additionally investigated subjective value representation as reported by others^68–70^ with univariate statistical analysis based on general linear models (GLM)^142^. Unless noted otherwise, all following analysis steps are implemented in the SPM package (version 12.7771) run in MATLAB. To investigate whether subjective value representations change under acute stress, we first identified value correlations in areas like the medial prefrontal cortex (mPFC) and orbitofrontal cortex (OFC) as reported before (see^67^ for a review). To do so, functional runs were preprocessed as in the other analysis and (fMRIPrep 20.2.2^115^) aligned to MNI152NLin2009cAsym space and then smoothed with a 6 mm kernel. The model of the first level analysis contained as regressors two intercept predictors for the event of the first and second option presentation and three parametric modulators for the value estimate at the respective time point and the value difference (*chosen-unchose*n, at time point of second option presentation). The value estimates were either objective expected value (EV), classical subjective expected utility (EU^40^), or our PMC model derived perceived expected value measure (pEV). As confounds we included physiological regressors derived with the RETROICOR method^139^ via the TAPAS software collection^140^ plus the 6 standard motion parameters and three low frequency noise parameters derived from preprocessing (fMRIPrep 20.2.2^115^). To identify potential subjective value representations, we took first-level contrasts of the respective regressors into a one-sample t-test at the second level. SPM illustrations were created with nilearn.plotting functions and cluster-level inference with nilearn.glm.cluster_level_inference, an implementation of All-Resolutions Inference^143^. ARI is a non-parametric thresholding procedure^143^ that mitigates the recently reported inflated false positive rates, due to the assumption specific functional forms of spatial autocorrelation, as in Gaussian Random Field Theory (GRF)^144,145^. We used a cluster forming threshold of p < 0.001 (z > 3.1) and the true positive rate to alpha=0.05.

To test the effects of stress, we ran two additional 2nd level analyses for the intercept and the parametric modulatory (pmod) SPMs of the second value (/option - for which we found the value correlation in the previous 2nd level analysis). Here, we included the contrast from both sessions and both groups and specified a design matrix that comprised a main, a session, a group and a sessionxgroup interaction regressors plus subject wise regressors (run via nilearn.glm.second_level.SecondLevelModel). With the sessionxgroup interaction contrast, we then performed cluster level inference as before^143^ plus correcting for a small volume with a mPFC mask (MarsAtlas^71^).

## Declaration of interests

The authors declare no competing interests.

## Acknowledgements

We are grateful to C. Schnyder, K. Treiber and M. Moisa at the Zurich Center for Neuroeconomics for their excellent assistance in recruitment and participant facilitation. M.R., G.d.H., G.A., S.B., and C.C.R. were supported by the University Research Priority Program (URPP) ‘Adaptive Brain Circuits in Development and Learning’ (AdaBD) at the University of Zurich.C.C.R. was also funded by the Swiss National Science Foundation SNSF (grant no. 100019L-173248). G.d.H. was also funded by the Dutch Research Council NWO (Rubicon grant no. 019.183SG.017/8O3B) and the University of Zurich (Forschungskredit grant no. K-33153-02-01) C.L. received funding connected to the research by the Austrian Science Fund “Effects of Acute Stress on Social Behavior” (10.55776/I3381), “Neuronal circuits in health and disease”: COE16 [10.55776/COE16], and “The role of incentives and uncertainty in prosocial behavior” [10.55776/PAT1936023]. P.F. is supported by a Marie Skłodowska-Curie Postdoctoral Fellowship from the European Commission (101107160).

## Footnotes

1 https://github.com/ruffgroup/bauer/

2 We assumed that if prior expectations on a plausible number range becomes systematically influenced by stress, it should affect the priors of both payoff magnitudes equally in log-space,as they share the same neural substrate of number processing.

## Notes

### Competing Interest Statement

The authors have declared no competing interest.

## References

1. Mykyta, L. Work Conditions and Serious Psychological Distress Among Working Adults Aged 18–64: United States, 2021. (2023) doi:10.15620/cdc:126566.

2. Golden, L. & Kim, J. Irregular work shifts, work schedule flexibility and associations with work-family conflict and work stress in the US. Work-Life Balance in the Modern Workplace. Interdisciplinary Perspectives from Work-Family Research, Law and Policy (2017).

3. Stress in America^TM^ 2020: A National Mental Health Crisis. https://www.apa.org/news/press/releases/stress/2020/report-october.

4. Jaremka, L. M. et al. Loneliness Promotes Inflammation During Acute Stress. Psychol. Sci. 24, 1089–1097 (2012).

5. Mohler-Kuo, M., Dzemaili, S., Foster, S., Werlen, L. & Walitza, S. Stress and Mental Health among Children/Adolescents, Their Parents, and Young Adults during the First COVID-19 Lockdown in Switzerland. Int. J. Environ. Res. Public Heal. 18, 4668 (2021).

6. Steptoe, A. & Kivimäki, M. Stress and cardiovascular disease. Nat. Rev. Cardiol. 9, 360–370 (2012).

7. Mofatteh, M., Kingdom, L. C., University of Oxford, Turl Street, Oxford OX1 3DR, United, Kingdom, M. C., University of Oxford, Merton Street, Oxford OX1 4DJ, United & Kingdom, S. W. D. S. of P., University of Oxford, South Parks Road, Oxford OX1 3RE, United. Risk factors associated with stress, anxiety, and depression among university undergraduate students. AIMS Public Heal. 8, 36–65 (2021).

8. Goeders, N. E. The impact of stress on addiction. Eur. Neuropsychopharmacol. 13, 435–441 (2003).

9. Coman, G. J., Burrows, G. D. & Evans, B. J. Stress and Anxiety as Factors in the Onset of Problem Gambling: Implications for Treatment. Stress Med. 13, 235–244 (1997).

10. Haushofer, J. & Fehr, E. On the psychology of poverty. Science 344, 862–867 (2014).

11. Cueva, C. et al. Cortisol and testosterone increase financial risk taking and may destabilize markets. Sci. Rep. 5, 11206 (2015).

12. Rabkin, J. G. & Struening, E. L. Live Events, Stress, and Illness. Science 194, 1013–1020 (1976).

13. Buckert, M., Schwieren, C., Kudielka, B. M. & Fiebach, C. J. Acute stress affects risk taking but not ambiguity aversion. Front. Neurosci. 8, 82 (2014).

14. Helversen, B. von & Rieskamp, J. Stress‐related changes in financial risk taking: Considering joint effects of cortisol and affect. Psychophysiology 57, e13560 (2020).

15. Porcelli, A. J. & Delgado, M. R. Acute Stress Modulates Risk Taking in Financial Decision Making. Psychol. Sci. 20, 278–283 (2008).

16. Yamakawa, K., Ohira, H., Matsunaga, M. & Isowa, T. Prolonged Effects of Acute Stress on Decision-Making under Risk: A Human Psychophysiological Study. Front. Hum. Neurosci. 10, 444 (2016).

17. Parslow, E. & Rose, J. Stress and risk — Preferences versus noise. Judgm. Decis. Mak. 17, 883–936 (2022).

18. Sokol-Hessner, P., Raio, C. M., Gottesman, S. P., Lackovic, S. F. & Phelps, E. A. Acute stress does not affect risky monetary decision-making. Neurobiol. Stress 5, 19–25 (2016).

19. Olschewski, S., Rieskamp, J. & Scheibehenne, B. Taxing Cognitive Capacities Reduces Choice Consistency Rather Than Preference: A Model-Based Test. J Exp Psychology Gen 147, 462–484 (2018).

20. Olschewski, S. & Rieskamp, J. Distinguishing three effects of time pressure on risk taking: Choice consistency, risk preference, and strategy selection. J. Behav. Decis. Mak. 34, 541–554 (2021).

21. Epel, E. S. et al. More than a feeling: A unified view of stress measurement for population science. Front. Neuroendocr. 49, 146–169 (2018).

22. Porcelli, A. J. & Delgado, M. R. Acute Stress Modulates Risk Taking in Financial Decision Making. Psychol. Sci. 20, 278–283 (2008).

23. Parslow, E. & Rose, J. Stress and risk — Preferences versus noise. Judgm. Decis. Mak. 17, 883–936 (2022).

24. Pabst, S., Brand, M. & Wolf, O. T. Stress and decision making: A few minutes make all the difference. Behav. Brain Res. 250, 39–45 (2013).

25. Helversen, B. von & Rieskamp, J. Stress‐related changes in financial risk taking: Considering joint effects of cortisol and affect. Psychophysiology 57, e13560 (2020).

26. Leith, K. P. & Baumeister, R. F. Why do bad moods increase self-defeating behavior? Emotion, risk tasking, and self-regulation. J. Pers. Soc. Psychol. 71, 1250–1267 (1996).

27. Preston, S. D., Buchanan, T. W., Stansfield, R. B. & Bechara, A. Effects of Anticipatory Stress on Decision Making in a Gambling Task. Behav. Neurosci. 121, 257–263 (2007).

28. Lighthall, N. R., Mather, M. & Gorlick, M. A. Acute Stress Increases Sex Differences in Risk Seeking in the Balloon Analogue Risk Task. PLoS ONE 4, e6002 (2009).

29. Yamakawa, K., Ohira, H., Matsunaga, M. & Isowa, T. Prolonged Effects of Acute Stress on Decision-Making under Risk: A Human Psychophysiological Study. Front. Hum. Neurosci. 10, 444 (2016).

30. Khaw, M. W., Li, Z. & Woodford, M. Cognitive Imprecision and Small-Stakes Risk Aversion. Rev. Econ. Stud. 88, 1979–2013 (2020).

31. Khaw, M. W., Li, Z. & Woodford, M. Risk Aversion as a Perceptual Bias. (2017) doi:10.3386/w23294.

32. Polanía, R., Burdakov, D. & Hare, T. A. Rationality, preferences, and emotions with biological constraints: it all starts from our senses. Trends Cogn. Sci. 28, 264–277 (2024).

33. Barretto-García, M. et al. Individual risk attitudes arise from noise in neurocognitive magnitude representations. Nat. Hum. Behav. 7, 1551–1567 (2023).

34. Urai, A. E., Braun, A. & Donner, T. H. Pupil-linked arousal is driven by decision uncertainty and alters serial choice bias. 8, (2017).

35. Gee, J. W. de et al. Pupil-linked phasic arousal predicts a reduction of choice bias across species and decision domains. eLife 9, e54014 (2020).

36. Hollander, G. de, Grueschow, M., Hennel, F. & Ruff, C. C. Rapid Changes in Risk Preferences Originate from Bayesian Inference on Parietal Magnitude Representations. bioRxiv 2024.08.23.609296 (2024) doi:10.1101/2024.08.23.609296.

37. Khaw, M. W., Li, Z. & Woodford, M. Cognitive Imprecision and Small-Stakes Risk Aversion. Rev Econ Stud 88, 1979–2013 (2020).

38. de Hollander, G., Grueschow, M., Hennel, F. & Ruff, C. C. Rapid Changes in Risk Preferences Originate from Bayesian Inference on Parietal Magnitude Representations. bioRxiv 2024.08.23.609296 (2024) doi:10.1101/2024.08.23.609296.

39. Kahneman, D. & Tversky, A. Prospect Theory: An Analysis of Decision under Risk. Econometrica 47, 263 (1979).

40. Neumann, J. V. & Morgenstern, O. Theory of Games and Economic Behavior, 2nd Rev. Ed. (Princeton University Press, Princeton, NJ, US, 1947).

41. McFadden & D. Conditional Logit Analysis of Qualitative Choice Behavior. Frontiers in Econometrics 105–142 (1974).

42. Heng, J. A., Woodford, M. & Polania, R. Efficient sampling and noisy decisions. eLife 9, e54962 (2020).

43. Prat-Carrabin, A. & Woodford, M. Efficient coding of numbers explains decision bias and noise. Nat. Hum. Behav. 6, 1142–1152 (2022).

44. Feigenson, L., Dehaene, S. & Spelke, E. Core systems of number. Trends Cogn. Sci. 8, 307–314 (2004).

45. Nieder, A. & Dehaene, S. Representation of Number in the Brain. Annu. Rev. Neurosci. 32, 185–208 (2009).

46. Bueti, D. & Walsh, V. The parietal cortex and the representation of time, space, number and other magnitudes. Philos. Trans. R. Soc. B: Biol. Sci. 364, 1831–1840 (2009).

47. Nieder, A. & Miller, E. K. A parieto-frontal network for visual numerical information in the monkey. Proc. Natl. Acad. Sci. 101, 7457–7462 (2004).

48. Harvey, B. M., Klein, B. P., Petridou, N. & Dumoulin, S. O. Topographic Representation of Numerosity in the Human Parietal Cortex. Science 341, 1123–1126 (2013).

49. Cai, Y. et al. Topographic numerosity maps cover subitizing and estimation ranges. Nat. Commun. 12, 3374 (2021).

50. Lasne, G., Piazza, M., Dehaene, S., Kleinschmidt, A. & Eger, E. Discriminability of numerosity-evoked fMRI activity patterns in human intra-parietal cortex reflects behavioral numerical acuity. Cortex 114, 90–101 (2019).

51. Woodford, M. Modeling Imprecision in Perception, Valuation, and Choice. Annu. Rev. Econ. 12, 1–23 (2020).

52. Olschewski, S., Rieskamp, J. & Hertwig, R. The link between cognitive abilities and risk preference depends on measurement. Sci. Rep. 13, 21151 (2023).

53. Bhatia, S. & Loomes, G. Noisy Preferences in Risky Choice: A Cautionary Note. Psychol. Rev. 124, 678–687 (2017).

54. Starcke, K., Wolf, O. T., Markowitsch, H. J. & Brand, M. Anticipatory Stress Influences Decision Making Under Explicit Risk Conditions. Behav. Neurosci. 122, 1352–1360 (2008).

55. Starcke, K. & Brand, M. Decision making under stress: A selective review. Neurosci. Biobehav. Rev. 36, 1228–1248 (2012).

56. Degroote, C. et al. Acute Stress Improves Concentration Performance. Exp. Psychol. 67, 88–98 (2020).

57. Globig, L. K., Witte, K., Feng, G. & Sharot, T. Under Threat, Weaker Evidence Is Required to Reach Undesirable Conclusions. J. Neurosci. 41, 6502–6510 (2021).

58. Bowers, J. S. & Davis, C. J. Bayesian Just-So Stories in Psychology and Neuroscience. Psychol. Bull. 138, 389–414 (2012).

59. Schildberg-Hörisch, H. Are Risk Preferences Stable? J. Econ. Perspect. 32, 135–154 (2018).

60. Dedovic, K. et al. The Montreal Imaging Stress Task: using functional imaging to investigate the effects of perceiving and processing psychosocial stress in the human brain. J. psychiatry Neurosci. : JPN 30, 319–25 (2005).

61. Shaffer, F. & Ginsberg, J. P. An Overview of Heart Rate Variability Metrics and Norms. Front. Public Heal. 5, 258 (2017).

62. Castaldo, R. et al. Acute mental stress assessment via short term HRV analysis in healthy adults: A systematic review with meta-analysis. Biomed. Signal Process. Control 18, 370–377 (2015).

63. Visnovcova, Z. et al. Complexity and time asymmetry of heart rate variability are altered in acute mental stress. Physiol. Meas. 35, 1319–1334 (2014).

64. Forbes, P. A. et al. Acute stress reduces effortful prosocial behaviour. eLife 12, RP87271 (2024).

65. Hollander, G. de, Moisa, M. & Ruff, C. C. Risk preferences causally rely on parietal magnitude representations: Evidence from combined TMS-fMRI. bioRxiv 2025.01.13.632678 (2025) doi:10.1101/2025.01.13.632678.

66. Vehtari, A., Gelman, A. & Gabry, J. Practical Bayesian model evaluation using leave-one-out cross-validation and WAIC. Stat Comput 27, 1413–1432 (2017).

67. Levy, D. J. & Glimcher, P. W. The root of all value: a neural common currency for choice. Curr. Opin. Neurobiol. 22, 1027–1038 (2012).

68. Kable, J. W. & Glimcher, P. W. The neural correlates of subjective value during intertemporal choice. Nat. Neurosci. 10, 1625–1633 (2007).

69. Levy, I., Snell, J., Nelson, A. J., Rustichini, A. & Glimcher, P. W. Neural Representation of Subjective Value Under Risk and Ambiguity. J. Neurophysiol. 103, 1036–1047 (2010).

70. Peters, J. & Büchel, C. Overlapping and Distinct Neural Systems Code for Subjective Value during Intertemporal and Risky Decision Making. J. Neurosci. 29, 15727–15734 (2009).

71. Auzias, G., Coulon, O. & Brovelli, A. MarsAtlas: A cortical parcellation atlas for functional mapping. Hum. Brain Mapp. 37, 1573–1592 (2016).

72. Buckert, M., Schwieren, C., Kudielka, B. M. & Fiebach, C. J. Acute stress affects risk taking but not ambiguity aversion. Front. Neurosci. 8, 82 (2014).

73. Bendahan, S. et al. Acute stress alters individual risk taking in a time‐dependent manner and leads to anti‐social risk. Eur. J. Neurosci. 45, 877–885 (2017).

74. Kandasamy, N. et al. Cortisol shifts financial risk preferences. Proc. Natl. Acad. Sci. 111, 3608–3613 (2014).

75. Gee, J. W. de, Knapen, T. & Donner, T. H. Decision-related pupil dilation reflects upcoming choice and individual bias. Proc. Natl. Acad. Sci. 111, E618–E625 (2014).

76. Gee, J. W. de et al. Pupil-linked phasic arousal predicts a reduction of choice bias across species and decision domains. eLife 9, e54014 (2020).

77. Ulrich-Lai, Y. M. & Herman, J. P. Neural regulation of endocrine and autonomic stress responses. Nat. Rev. Neurosci. 10, 397–409 (2009).

78. Belli, F., Felisatti, A. & Fischer, M. H. “BreaThink”: breathing affects production and perception of quantities. Exp. Brain Res. 239, 2489–2499 (2021).

79. Meck, W. H. & Church, R. M. A mode control model of counting and timing processes. J. Exp. Psychol.: Anim. Behav. Process. 9, 320–334 (1983).

80. Hedger, K. van, Necka, E. A., Barakzai, A. K. & Norman, G. J. The influence of social stress on time perception and psychophysiological reactivity. Psychophysiology 54, 706–712 (2017).

81. Weber, E. U., Blais, A. & Betz, N. E. A domain‐specific risk‐attitude scale: measuring risk perceptions and risk behaviors. J. Behav. Decis. Mak. 15, 263–290 (2002).

82. Frey, R., Pedroni, A., Mata, R., Rieskamp, J. & Hertwig, R. Risk preference shares the psychometric structure of major psychological traits. Sci. Adv. 3, e1701381 (2017).

83. Pedroni, A. et al. The risk elicitation puzzle. Nat Hum Behav 1, 803–809 (2017).

84. Walker, E. Y. et al. Studying the neural representations of uncertainty. Nat. Neurosci. 1–11 (2023) doi:10.1038/s41593-023-01444-y.

85. Vilares, I., Howard, J. D., Fernandes, H. L., Gottfried, J. A. & Kording, K. P. Differential Representations of Prior and Likelihood Uncertainty in the Human Brain. Curr. Biol. 22, 1641–1648 (2012).

86. Ting, C.-C., Yu, C.-C., Maloney, L. T. & Wu, S.-W. Neural Mechanisms for Integrating Prior Knowledge and Likelihood in Value-Based Probabilistic Inference. J. Neurosci. 35, 1792–1805 (2015).

87. Meyniel, F. & Dehaene, S. Brain networks for confidence weighting and hierarchical inference during probabilistic learning. Proc. Natl. Acad. Sci. 114, E3859–E3868 (2017).

88. Ma, W. J., Beck, J. M., Latham, P. E. & Pouget, A. Bayesian inference with probabilistic population codes. Nat. Neurosci. 9, 1432–1438 (2006).

89. Anton-Erxleben, K. & Carrasco, M. Attentional enhancement of spatial resolution: linking behavioural and neurophysiological evidence. Nat. Rev. Neurosci. 14, 188–200 (2013).

90. Klein, B. P. et al. Cortical depth dependent population receptive field attraction by spatial attention in human V1. NeuroImage 176, 301–312 (2018).

91. Dumoulin, S. O., Fracasso, A., Zwaag, W. van der, Siero, J. C. W. & Petridou, N. Ultra-high field MRI: Advancing systems neuroscience towards mesoscopic human brain function. NeuroImage 168, 345–357 (2018).

92. Stephan, K. E. et al. Laminar fMRI and computational theories of brain function. Neuroimage 197, 699–706 (2019).

93. Klein, B. P., Harvey, B. M. & Dumoulin, S. O. Attraction of Position Preference by Spatial Attention throughout Human Visual Cortex. Neuron 84, 227–237 (2014).

94. van Es, D. M., Theeuwes, J. & Knapen, T. Spatial sampling in human visual cortex is modulated by both spatial and feature-based attention. Elife 7, e36928 (2018).

95. Spurrier, M. & Blaszczynski, A. Risk Perception in Gambling: A Systematic Review. J. Gambl. Stud. 30, 253–276 (2014).

96. Hartley, C. A. & Phelps, E. A. Anxiety and Decision-Making. Biol. Psychiatry 72, 113–118 (2012).

97. Kirschbaum, C., Kudielka, B. M., Gaab, J., Schommer, N. C. & Hellhammer, D. H. Impact of Gender, Menstrual Cycle Phase, and Oral Contraceptives on the Activity of the Hypothalamus-Pituitary-Adrenal Axis. Psychosom. Med. 61, 154–162 (1999).

98. Tomova, L. et al. Increased neural responses to empathy for pain might explain how acute stress increases prosociality. Soc. Cogn. Affect. Neurosci. 12, 401–408 (2017).

99. Pruessner, J. C., Kirschbaum, C., Meinlschmid, G. & Hellhammer, D. H. Two formulas for computation of the area under the curve represent measures of total hormone concentration versus time-dependent change. Psychoneuroendocrinology 28, 916–931 (2003).

100. Makowski, D. et al. NeuroKit2: A Python toolbox for neurophysiological signal processing. Behav. Res. Methods 53, 1689–1696 (2021).

101. Capretto, T. et al. Bambi: A simple interface for fitting Bayesian linear models in Python. arXiv (2020) doi:10.48550/arxiv.2012.10754.

102. Hollander, G. de, Renkert, M. F. & Ruff, C. C. Bauer: Bayesian Estimation of Perceptual, Numerical and Risky Judgements. (2024).

103. Patil, A., Huard, D. & Fonnesbeck, C. J. PyMC: Bayesian Stochastic Modelling in Python. J. Stat. Softw. 35, 1–81 (2010).

104. Hoffman, M. D. & Gelman, A. The No-U-Turn Sampler: Adaptively Setting Path Lengths in Hamiltonian Monte Carlo. Journal of Machine Learning Research 15, 1593–1623 (2011).

105. Kruschke, J. K. Doing Bayesian Data Analysis: A Tutorial with R and BUGS. (Academic Press., Burlington, MA, 2011).

106. Olschewski, S. & Rieskamp, J. Distinguishing three effects of time pressure on risk taking: Choice consistency, risk preference, and strategy selection. J. Behav. Decis. Mak. 34, 541–554 (2021).

107. Capretto, T. et al. Bambi: A simple interface for fitting Bayesian linear models in Python. arXiv (2020) doi:10.48550/arxiv.2012.10754.

108. Garcia, M. B. et al. Individual risk attitudes arise from noise in neurocognitive magnitude representations. Biorxiv 2022.08.22.504413 (2022) doi:10.1101/2022.08.22.504413.

109. Hoffman, M. D. & Gelman, A. The No-U-Turn Sampler: Adaptively Setting Path Lengths in Hamiltonian Monte Carlo. Journal of Machine Learning Research 15, 1593–1623 (2011).

110. Patil, A., Huard, D. & Fonnesbeck, C. J. PyMC: Bayesian Stochastic Modelling in Python. J. Stat. Softw. 35, 1–81 (2010).

111. Betancourt, M. J. & Girolami, M. Hamiltonian Monte Carlo for Hierarchical Models. arXiv 79, 2–4 (2015).

112. Wiecki, T. V., Sofer, I. & Frank, M. J. HDDM: Hierarchical Bayesian estimation of the Drift-Diffusion Model in Python. Front Neuroinform 7, 14 (2013).

113. Gelman, A. Prior distributions for variance parameters in hierarchical models (comment on article by Browne and Draper). Bayesian Anal 1, 515–534 (2006).

114. Kruschke, J. K. Doing Bayesian Data Analysis: A Tutorial with R and BUGS. (Academic Press., Burlington, MA, 2011).

115. Esteban, O. et al. fMRIPrep: a robust preprocessing pipeline for functional MRI. Nat Methods 16, 111–116 (2019).

116. Gorgolewski, K. J. et al. Nipype. Software. Zenodo. 10.5281/zenodo596855, (2018).

117. Gorgolewski, K. J. et al. Nipype: A Flexible, Lightweight and Extensible Neuroimaging Data Processing Framework in Python. Frontiers Neuroinformatics 5, 13 (2011).

118. Tustison, N. J. et al. N4ITK: Improved N3 Bias Correction. IEEE Trans. Méd. Imaging 29, 1310–1320 (2010).

119. Avants, B. B., Tustison, N. & Song, G. Advanced normalization tools (ANTS). 2, (2009).

120. Zhang, Y., Brady, M. & Smith, S. Segmentation of Brain MR Images Through a Hidden Markov Random Field Model and the Expectation-Maximization Algorithm. IEEE Trans. Méd. Imaging 20, 45 (2001).

121. Reuter, M., Rosas, H. D. & Fischl, B. Highly accurate inverse consistent registration: A robust approach. NeuroImage 53, 1181–1196 (2010).

122. Dale, A. M., Fischl, B. & Sereno, M. I. Cortical Surface-Based Analysis I. Segmentation and Surface Reconstruction. Neuroimage 9, 179–194 (1999).

123. Klein, A. et al. Mindboggling morphometry of human brains. PLoS Comput. Biology 13, e1005350 (2017).

124. Fonov, V., Evans, A., McKinstry, R., Almli, C. & Collins, D. Unbiased nonlinear average age-appropriate brain templates from birth to adulthood. NeuroImage 47, S102 (2009).

125. Cox, R. W. & Hyde, J. S. Software tools for analysis and visualization of fMRI data. NMR Biomed. 10, 171–178 (1997).

126. Jenkinson, M., Bannister, P., Brady, M. & Smith, S. Improved optimization for the robust and accurate linear registration and motion correction of brain images. Neuroimage 17, 825–841 (2002).

127. Greve, D. N. & Fischl, B. Accurate and robust brain image alignment using boundary-based registration. 48, (2009).

128. Power, J. D. et al. Methods to detect, characterize, and remove motion artifact in resting state fMRI. NeuroImage 84, 320–341 (2014).

129. Behzadi, Y., Restom, K., Liau, J. & Liu, T. T. A component based noise correction method (CompCor) for BOLD and perfusion based fMRI. NeuroImage 37, 90–101 (2007).

130. Satterthwaite, T. D. et al. An improved framework for confound regression and filtering for control of motion artifact in the preprocessing of resting-state functional connectivity data. NeuroImage 64, 240–256 (2013).

131. Lanczos, C. Evaluation of Noisy Data. J. Soc. Ind. Appl. Math. Ser. B Numer. Anal. 1, 76–85 (1964).

132. Abraham, A. et al. Machine learning for neuroimaging with scikit-learn. Frontiers Neuroinformatics 8, 14 (2014).

133. van Bergen, R. S., Ma, W. J., Pratte, M. S. & Jehee, J. F. M. Sensory uncertainty decoded from visual cortex predicts behavior. Nat Neurosci 18, 1728 (2015).

134. Walker, E. Y., Cotton, R. J., Ma, W. J. & Tolias, A. S. A neural basis of probabilistic computation in visual cortex. Nat Neurosci 23, 122–129 (2020).

135. Harvey, B. M., Klein, B. P., Petridou, N. & Dumoulin, S. O. Topographic Representation of Numerosity in the Human Parietal Cortex. Science 341, 1123–1126 (2013).

136. Prince, J. S. et al. Improving the accuracy of single-trial fMRI response estimates using GLMsingle. Elife 11, e77599 (2022).

137. Mumford, J. A., Turner, B. O., Ashby, F. G. & Poldrack, R. A. Deconvolving BOLD activation in event-related designs for multivoxel pattern classification analyses. NeuroImage 59, 2636–2643 (2012).

138. Kay, K. N., Rokem, A., Winawer, J., Dougherty, R. F. & Wandell, B. A. GLMdenoise: a fast, automated technique for denoising task-based fMRI data. Front. Neurosci. 7, 247 (2013).

139. Glover, G. H., Li, T. & Ress, D. Image‐based method for retrospective correction of physiological motion effects in fMRI: RETROICOR. Magn. Reson. Med. 44, 162–167 (2000).

140. Frässle, S. et al. TAPAS: An Open-Source Software Package for Translational Neuromodeling and Computational Psychiatry. Front. Psychiatry 12, 680811 (2021).

141. Kingma, D. P. & Ba, J. Adam: A Method for Stochastic Optimization. arXiv (2014) doi:10.48550/arxiv.1412.6980.

142. Friston, K. J. et al. Statistical parametric maps in functional imaging: A general linear approach. Hum. Brain Mapp. 2, 189–210 (1994).

143. Rosenblatt, J. D., Finos, L., Weeda, W. D., Solari, A. & Goeman, J. J. All-Resolutions Inference for brain imaging. NeuroImage 181, 786–796 (2018).

144. Eklund, A., Nichols, T. E. & Knutsson, H. Cluster failure: Why fMRI inferences for spatial extent have inflated false-positive rates. Proc. Natl. Acad. Sci. 113, 7900–7905 (2016).

145. Kempen, M. van. Validating the All-Resolutions Inference method for analyzing fMRI and building a comprehensive app for the end users. Master’s thesis, Leiden University, Leiden University Repository.

146. Paul, J. M., Ackooij, M. van, Cate, T. C. ten & Harvey, B. M. Numerosity tuning in human association cortices and local image contrast representations in early visual cortex. Nat. Commun. 13, 1340 (2022).

