## Supplements for "Acute stress reduces risk-aversion by changing magnitude perception"

### Supplementary Material

Supplementary Table 1: Parameter estimates (mean and confidence interval (CI)) for all group/session-pairs. Note that the means (mu) and the standard deviations (sd) are in log-space.

| Parameter | Group/Session | Mean | CI Lower | CI Upper |
|---:|---:|---:|---:|---:|
| risky_prior_mu | Group 0, Session 1 | 3.291343 | 2.993922 | 3.577278 |
| | Group 0, Session 2 | 3.171644 | 2.945387 | 3.391746 |
| | Group 1, Session 1 | 2.771165 | 2.411254 | 3.147748 |
| | Group 1, Session 2 | 3.158895 | 2.803672 | 3.500723 |
| safe_prior_mu | Group 0, Session 1 | 3.201961 | 2.713737 | 3.776340 |
| | Group 0, Session 2 | 3.082262 | 2.638719 | 3.632946 |
| | Group 1, Session 1 | 2.681782 | 2.116092 | 3.292029 |
| | Group 1, Session 2 | 3.069513 | 2.526623 | 3.686832 |
| n1_evidence_sd | Group 0, Session 1 | 0.283917 | 0.229173 | 0.349889 |
| | Group 0, Session 2 | 0.252987 | 0.222020 | 0.287219 |
| | Group 1, Session 1 | 0.298745 | 0.229232 | 0.381224 |
| | Group 1, Session 2 | 0.283775 | 0.243687 | 0.328330 |
| n2_evidence_sd | Group 0, Session 1 | 0.198643 | 0.161072 | 0.244184 |
| | Group 0, Session 2 | 0.192698 | 0.168315 | 0.218998 |
| | Group 1, Session 1 | 0.215563 | 0.167006 | 0.273226 |
| | Group 1, Session 2 | 0.204021 | 0.172681 | 0.236092 |
| risky_prior_sd | Group 0, Session 1 | 0.344085 | 0.239295 | 0.474368 |
| | Group 0, Session 2 | 0.367670 | 0.278464 | 0.475519 |
| | Group 1, Session 1 | 0.367770 | 0.245238 | 0.520930 |
| | Group 1, Session 2 | 0.367460 | 0.269576 | 0.490742 |
| safe_prior_sd | Group 0, Session 1 | 0.415288 | 0.242060 | 0.664322 |
| | Group 0, Session 2 | 0.601553 | 0.392841 | 0.886696 |
| | Group 1, Session 1 | 0.523857 | 0.274750 | 0.859193 |
| | Group 1, Session 2 | 0.678107 | 0.423307 | 1.051189 |

Supplementary Table 2: Effects of session and stress (session:group) on the widths of the risky and safe prior (distribution characterizing properties of the posterior chains of the regressors with $p_{bayesian}$ being the probability mass that is larger/smaller than zero)

| Parameter | Regressor | $p_{bayesian}$ | Mean | CI Lower | CI Upper |
|---|---|---|---|---|---|
| risky_prior_sd | session | 0.768 | 0.084970 | -0.137659 | 0.314362 |
| | session:group | 0.314 | -0.079264 | -0.395347 | 0.253085 |
| safe_prior_sd | session | 0.996 | 0.482564 | 0.118309 | 0.936636 |
| | session:group | 0.337 | -0.125639 | -0.734145 | 0.479683 |

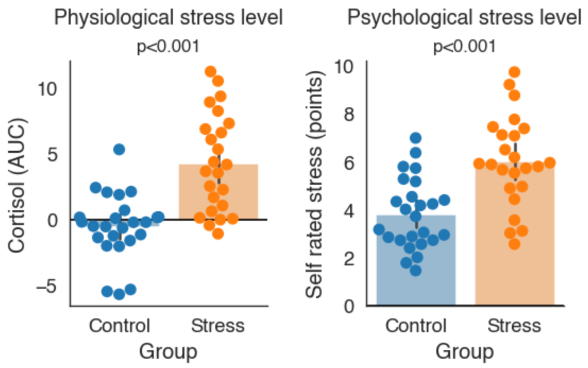


Supplementary Figure 1: Subject wise mean (over whole experiment of session two) stress level distributions per group, evaluated with physiological (AUC) and psychological measures (questionnaire). Test statistics in main text.

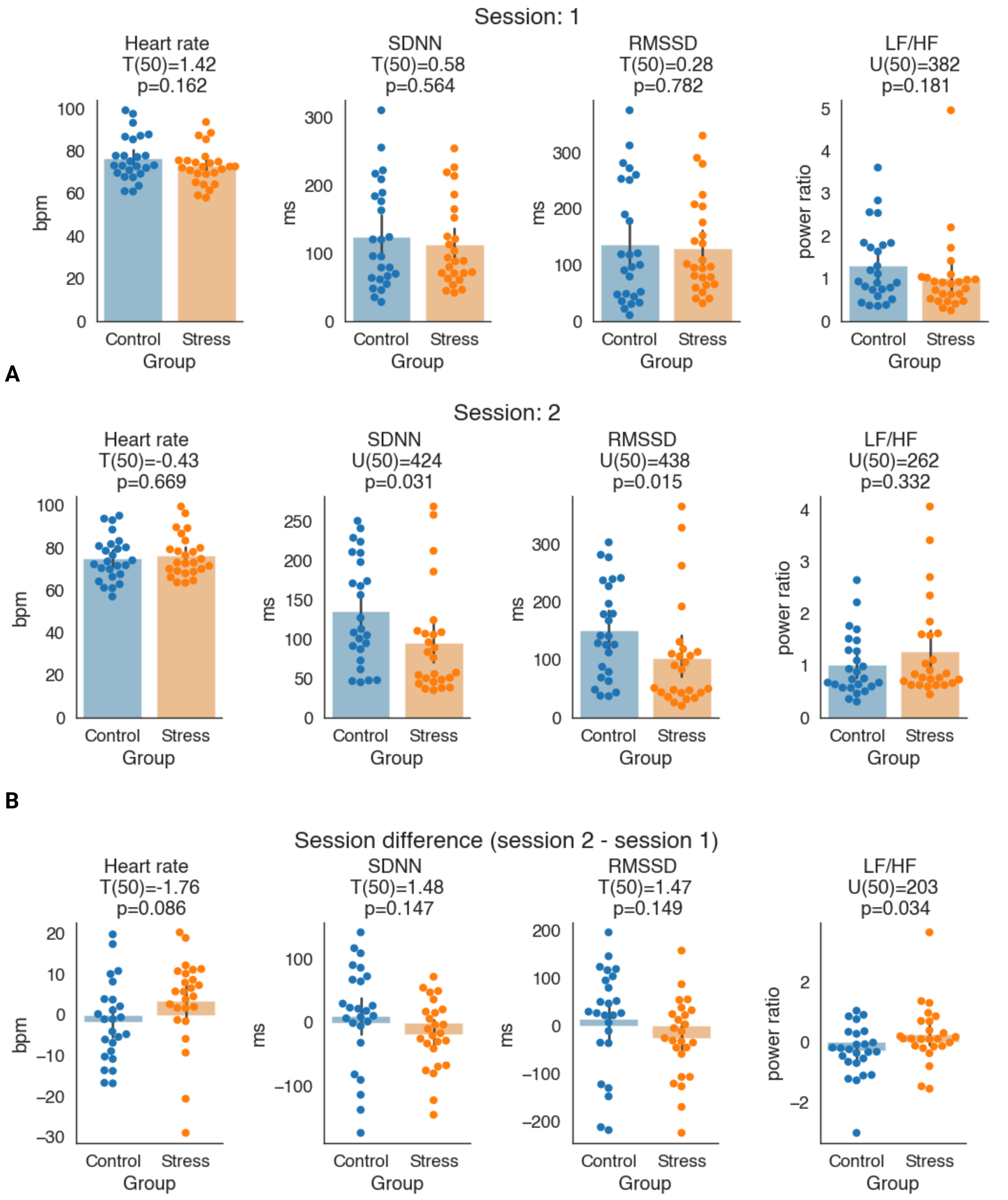


Supplementary Figure 2: Between groups comparison of subject wise heart rate related measures. A) Average session value B) Session differences. Between group test statistics given for each comparison, for normally distributed values as TTest and for non normally distributed as Mann-Whitney U test. Heart rate - in beats per minute (bpm); SDNN - Standard deviation of NN-intervals - in milliseconds (ms); RMSDD - Root Mean Square of Successive Differences - in milliseconds (ms); LF/HF - low frequency high frequency heart rate variability ratio - as power ratio.

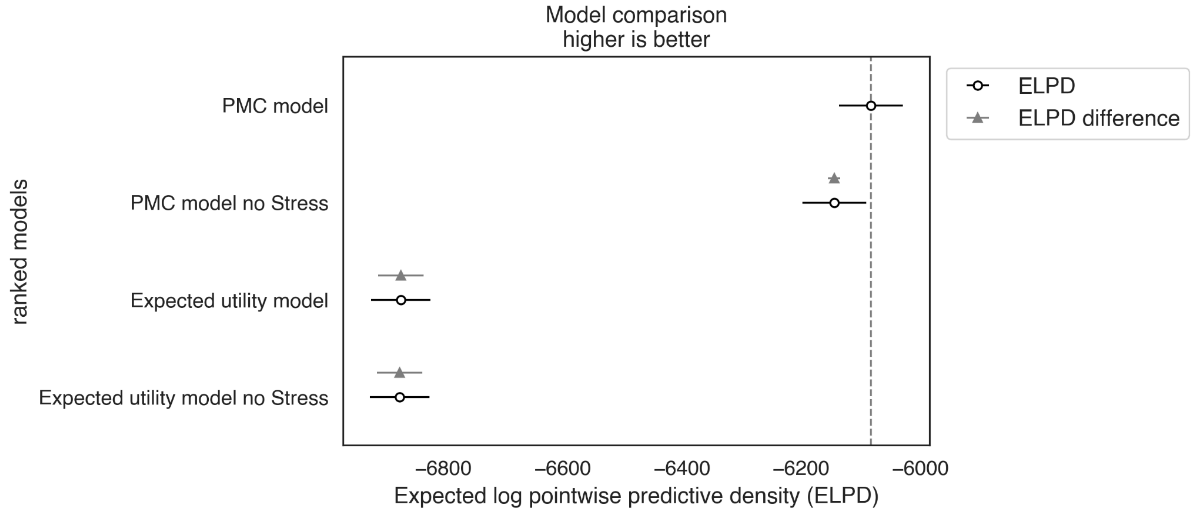


Supplementary Figure 3: Model comparison. We compared models with the expected log-predictive density (ELPD) as a measure of out-of-sample performance, which estimates the likelihood of unseen data given the model parameters[66]. It shows that a) our PMC model outperforms classical Expected Utility (EU) models, and b) models that comprise the stress regressor (as a session:group interaction effect) are better than their version without it.

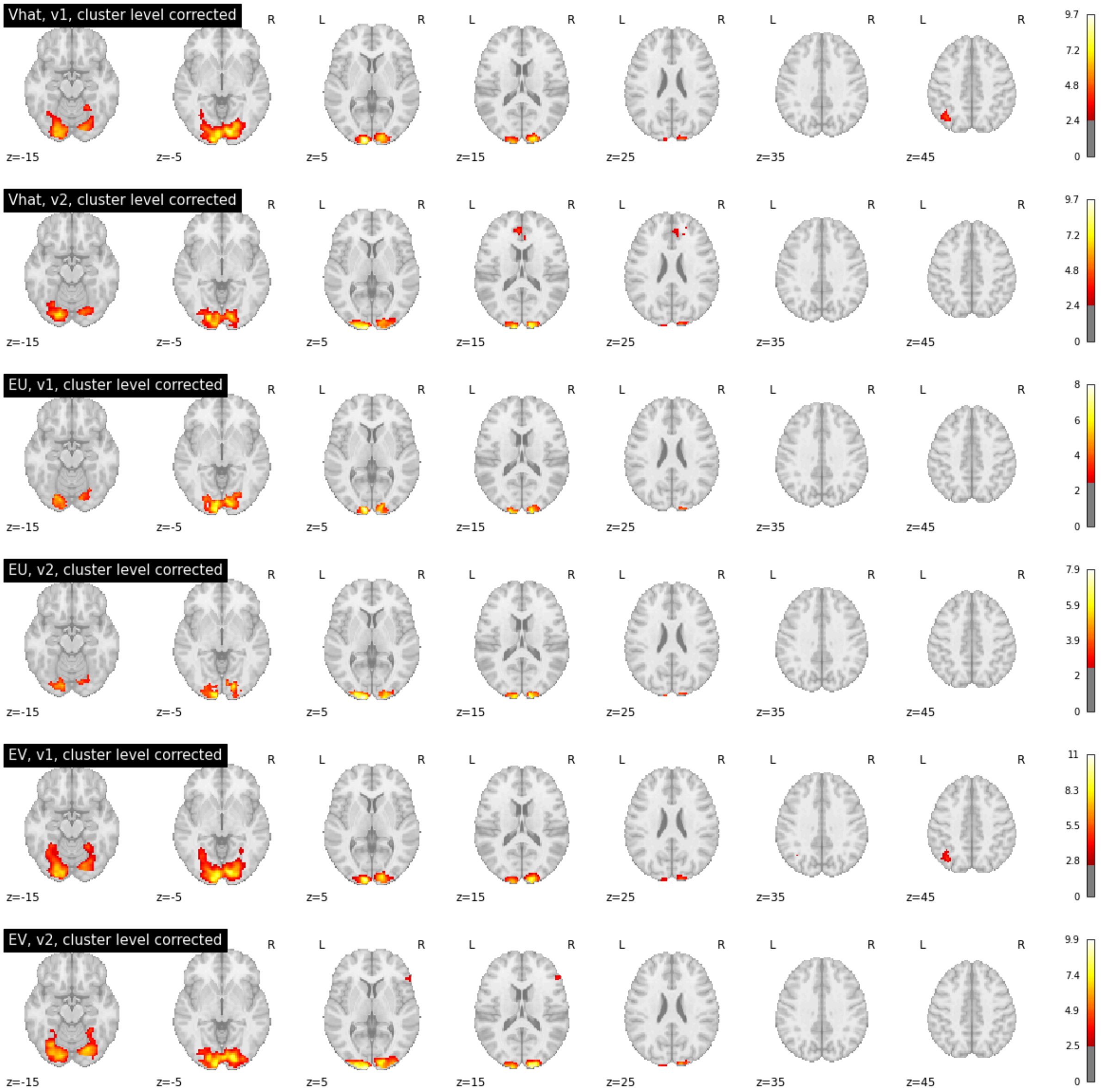


Supplementary Figure 4: Horizontal cross sections of statistical parametric maps from correlations (parametric modulator - pmod) with values from first (v1) and second (v2) option presentation, with different value estimates: Vhat = pEV= perceptual model; EU = classical expected utility model, EV = objective mathematical expectation. Visual cortices showed a reliable correlation to value, as one would expect given its monotonic response to spatial frequency that is tightly coupled to numerosity and hence value[146]. Only for Vhat-v2 there was a cluster in prefrontal values regions[68–70] that survived statistical thresholding (cluster forming threshold: $p<0.001$; family-wise error rate: $\alpha < 0.05$). Cluster details are reported in the tables below.

pEV - option 1

| Cluster ID | X | Y | Z | Peak Stat | Cluster Size (mm3) |
|---|---|---|---|---|---|
| 1 | -11.5 | -100.0 | 5.5 | 9.665699 | 36450 |
| 2 | -41.5 | -65.0 | 50.5 | 4.811005 | 2400 |

pEV - option 2

| Cluster ID | X | Y | Z | Peak Stat | Cluster Size (mm3) |
|---|---|---|---|---|---|
| 1 | -11.5 | -102.5 | 8.5 | 9.652147 | 27975 |
| 2 | -4.0 | 40.0 | 17.5 | 4.870433 | 2400 |

EU - option 1

| Cluster ID | X | Y | Z | Peak Stat | Cluster Size (mm3) |
|---|---|---|---|---|---|
| 1 | -11.5 | -100.0 | -0.5 | 7.9911 | 20737 |

EU - option 2

| Cluster ID | X | Y | Z | Peak Stat | Cluster Size (mm3) |
|---|---|---|---|---|---|
| 1 | -11.5 | -100.0 | -3.5 | 7.8726597 | 10050 |
| 2 | 13.5 | -102.5 | 11.5 | 6.756739 | 6431 |

EV - option 1

| Cluster ID | X | Y | Z | Peak Stat | Cluster Size (mm3) |
|---|---|---|---|---|---|
| 1 | -11.5 | -100.0 | 2.5 | 11.007339 | 46537 |
| 2 | -41.5 | -65.0 | 50.5 | 5.097252 | 3131 |

EV - option 2

| Cluster ID | X | Y | Z | Peak Stat | Cluster Size (mm3) |
|---|---|---|---|---|---|
| 1 | -14.0 | -105.0 | 8.5 | 9.89545 | 42468 |
| 2 | 51.0 | 30.0 | 11.5 | 5.079841 | 768 |
